# 3′ Plasma Membrane Phosphoinositides Sustain ROS Production at Persistent Non-canonical Phagocytic Cups that Promote Macrophage Control of *Aspergillus fumigatus* Hyphae

**DOI:** 10.64898/2026.09.14.751521

**Authors:** Serene Moussaoui, Maria Cecilia Gimenez, Amanda Paixao, Melie Boulianne, Micaela Cabrini, Amir Kanbar, Abdumalik Okhunjonov, Carlos Arnillas Merino, Mauricio R. Terebiznik

## Abstract

*Aspergillus fumigatus* is the leading cause of aspergillosis in immunocompromised individuals. Because *A. fumigatus* hyphae rapidly outgrow macrophages, they are generally thought to evade macrophage-mediated control once they exceed the size limits of phagocytosis, enabling invasive growth within host tissues. Here, we show that macrophages circumvent this limitation by capturing hyphal tips within persistent tubular phagocytic cups. Rather than resolving into complete phagosomes or disengaging from their targets, these long-lived open compartments maintain key features of phagocytic cups while progressively acquiring characteristics of maturing phagosomes. Their persistence sustains localized NOX2-dependent reactive oxygen species production, leading to hyphal-tip damage and restriction of fungal growth. Sustained PI(3,4)P_2_ signaling downstream of class I PI3K promotes prolonged NOX2 activity and is required for efficient hyphal restriction. Together, our findings reveal a previously unrecognized antifungal mechanism in which persistent phagocytic cups transform incomplete phagocytosis into sustained antimicrobial activity, extending macrophage control beyond the physical limits of engulfment.

## Introduction

*Aspergillus fumigatus* (*A. fumigatus*) is a ubiquitous environmental mold and a major opportunistic fungal pathogen responsible for substantial morbidity and mortality among immunocompromised individuals ^1^. In healthy individuals, inhaled conidia are cleared by innate immune mechanisms, preventing the generation of hyphae and disease. In contrast, impaired host immunity allows conidial germination into hyphae that rapidly invade host tissues ^2^. *A. fumigatus* causes a spectrum of diseases whose manifestations are determined by host immune competence and underlying lung pathology, ranging from airway colonization and chronic allergic bronchopulmonary aspergillosis to invasive pulmonary aspergillosis ^3^. Reflecting its growing clinical and public health impact, *A. fumigatus* was designated a critical-priority fungal pathogen by the World Health Organization ^4^.

The prevailing paradigm of host defense against *A. fumigatus* centers on a sequential macrophage-to-neutrophil response ^1^. Within the alveolar compartment, alveolar macrophages constitute the first line of innate cellular defense by phagocytosing inhaled conidia. Following phagocytosis, conidia are trafficked to phagolysosomes where they are inactivated by the combined action of reactive oxygen and nitrogen species, hydrolytic enzymes, and nutrient deprivation ^5,6^. However, a subset of internalized conidia can survive intracellularly by adapting to the phagosomal environment and/or disrupting phagolysosomal maturation ^7–9^. Under conditions of impaired immune control, these surviving conidia may subsequently germinate. As *A. fumigatus* conidia germinate into hyphae, host antifungal defense transitions from a predominantly macrophage-mediated phagocytosis to neutrophil-driven extracellular antifungal activity^1^. This is believed to respond to fungal morphogenesis; whereas conidia are readily engulfed and killed by alveolar macrophages, hyphae rapidly elongate and branch, reaching dimensions that exceed the limits of efficient phagocytic uptake, thereby necessitating extracellular mechanisms for fungicidal defenses^1^. Consequently, recruited neutrophils emerge as the dominant effector cells against hyphae, restricting fungal growth through the production of reactive oxygen species (ROS), release of antimicrobial granule contents and the formation of neutrophil extracellular traps (NETs) ^1,10^.

Although the sequential macrophage-to-neutrophil response paradigm is widely accepted, accumulating evidence suggests it may underestimate macrophage contributions to defense against fungal hyphae ^11–14^. Macrophages possess a remarkable phagocytic capacity and can engulf targets whose dimensions approach or even exceed those of the phagocyte itself ^15^. Moreover, completion of phagocytosis is not dictated by target size alone, but is strongly influenced by membrane availability, target geometry and orientation, and engulfment mechanics ^16–18^. Fungal hyphae represent quintessential high-aspect-ratio targets, and macrophages have been shown to sustain long-lasting phagocytic cup interfaces during interactions with similarly elongated targets, including filamentous *Legionella pneumophila* (*L pneumophila*)*, Candida albicans* (*C. albicans*) hyphae, and engineered abiotic filaments ^12,19–23^. These persistent phagocytic structures support progressive engulfment and eventual internalization, underscoring the remarkable plasticity of macrophage phagocytosis, therefore suggesting that macrophages might play a role in the containment of fungal hyphae. Indeed, although prolonged neutropenia is a major risk factor for invasive aspergillosis, it is not sufficient to explain susceptibility as invasive disease develops only in a subset of neutropenic patients ^24–27^. Moreover, experimental depletion of alveolar macrophages and impairment of macrophage antifungal signaling pathways have been associated with reduced fungal clearance and increased susceptibility to pulmonary aspergillosis ^28^. Together, this evidence suggests that macrophage-mediated protection against *A. fumigatus* may extend beyond the initial control of inhaled conidia and contribute to restrict hyphal growth and invasion following germination.

Thus, to determine whether macrophages contribute directly to the control of *A. fumigatus* hyphal growth, we examined how these cells interact with fungal filaments that exceed their engulfment capacity. Here, we show that macrophages adapt their phagocytic machinery to form persistent tubular phagocytic cups (tPCs) which are hybrid compartments that retain features of both phagocytic cups and phagosomes. By holding on hyphal tips, the sites of polarized growth essential for fungal elongation, tPCs induce localized damage that restricts further hyphal extension. We found that this antifungal activity is sustained by prolonged NOX2-mediated ROS production and is associated with persistent enrichment of the phosphoinositide phosphatidylinositol 3,4-bisphosphate [PI(3,4)P_2_] at the cytosolic leaflet of the cup membrane, downstream of Class I PI3K activity. Together, these findings reveal a previously unrecognized macrophage adaptation and antimicrobial pathway that enables direct capture, containment, and damage of long hyphae, despite incomplete internalization. More broadly, our work expands the prevailing paradigm of antifungal immunity beyond the traditional division of labor in which macrophages eliminate conidia while neutrophils control hyphae.

## Results

### Macrophages grasp A. fumigatus hyphal tips within long-lasting tubular phagocytic cups

Macrophage microbicidal activity relies largely on phagocytosis and phagosomal degradation, raising the question of whether they can contend with fungal hyphae whose lengths preclude internalization ^29^. To investigate this, we challenged RAW 264.7 murine macrophage (RAW cells) with paraformaldehyde-fixed *A. fumigatus* hyphae. Fixation prevented fungal growth and secretion, enabling the effects of hyphal morphology to be examined independently of fungal-derived soluble factors and growth-associated mechanical forces that might otherwise impact phagocytosis. As shown in **Figure 1 A**, efficient macrophage binding to *A. fumigatus* hyphae depended on either IgG opsonization or expression of the β-1,3-glucan receptor Dectin-1A. In the absence of these interactions, binding was rare, indicating that phagocytic recognition is essential for efficient macrophage attachment to this target (**Figure 1 A**). Scanning Electron Microscopy (SEM) and time-lapse video microscopy revealed that macrophages attached to hyphal filaments and maintained these interactions for several hours (hrs), demonstrating the formation of persistent macrophage-hypha contacts (**Figure 1 B, Supplementary Videos 1-2**). Following attachment, macrophages spread extensively along the hyphal surface and adopted a morphology consistent with sustained engulfment of the target (**Figure 1 C**), despite the physical impossibility of completely internalizing the mature fungal filament. Instead, shorter fungal targets were efficiently internalized by macrophages up to a threshold length of approximately 80 μm and subsequently trafficked to phagosomes matured to acquire degradative capacity (**Supplementary Figure 1 A and B**).

**Figure 1.**
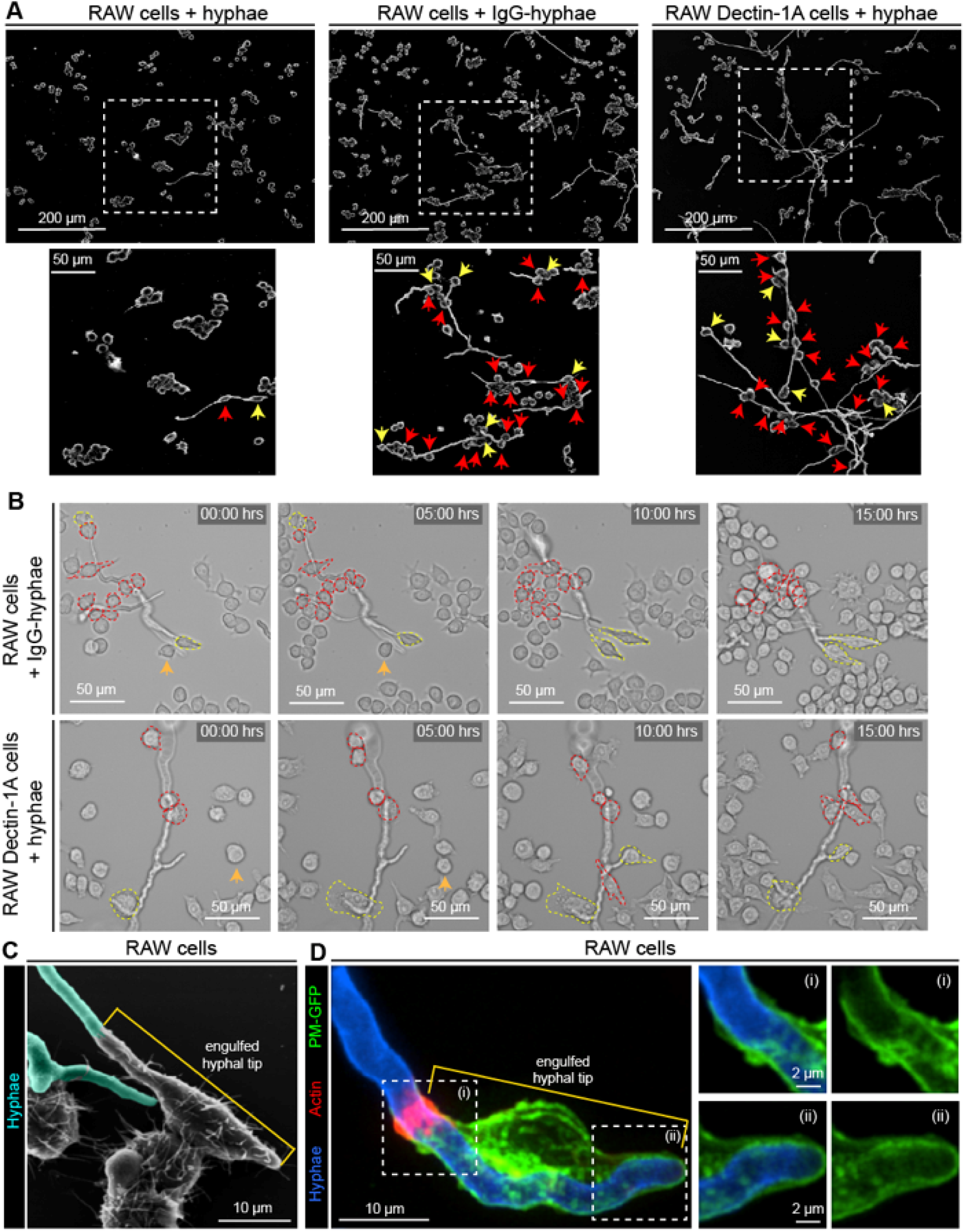
RAW macrophages sustain a phagocytic grasp on *A. fumigatus* hyphae. (**A-C**) Macrophage interactions with A. fumigatus hyphae. (**A**) Scanning electron microscopy (SEM) images of RAW and RAW Dectin-1A macrophages incubated for 3 hrs with paraformaldehyde-fixed non-opsonized hyphae (hyphae) or IgG-opsonized hyphae (IgG-hyphae). Yellow arrows indicate macrophages engaging hyphal tips, whereas red arrows indicate macrophages engaging along hyphal lengths. (**B**) RAW and RAW Dectin-1A macrophages were incubated with hyphae or IgG-hyphae and imaged by brightfield time-lapse microscopy for up to 15 hrs. Images show representative stills from Videos S1 and S2, respectively. Macrophages engaging hyphal tips are outlined with yellow dotted lines, whereas macrophages engaging along hyphal lengths are outlined with red dotted lines. Orange arrows indicate macrophages that are about to partake in hyphal tip engulfment. (**C**) SEM image of a RAW macrophage engulfing an IgG-hyphal tip. Hyphae are pseudocolored cyan. (**D**) tPC formation during hyphal engagement. RAW macrophages expressing PM-GFP (green) were incubated with IgG-hyphae (blue) for 1 hr, stained with phalloidin (red), and imaged by spinning-disk confocal microscopy (SDCM). Magnified views show the proximal (**i**) and distal (**ii**) regions of the tPC.

Hyphal tips are highly dynamic regions where cell wall synthesis and remodeling are concentrated to support apical growth ^30^. Thus, macrophage engagement of hyphal tips may selectively target the growth machinery responsible for filament elongation, potentially limiting fungal growth. Based on this rationale, we focused the present study on macrophage interactions with hyphal tips rather than lateral hyphal attachments. Confocal microscopy revealed that macrophages engulfed hyphal tips within plasma membrane invaginations crowned by a dense ring of filamentous actin (**Figure 1 D**). These structures are consistent with tubular phagocytic cups (tPCs), as their formation required phagocytic receptor engagement and they closely resembled those previously reported during the phagocytosis of filamentous bacteria and *Candida albicans* hyphae by our group and others ^12,19,23,31^.

### Hypha-holding tPCs feature phagosomal maturation hallmarks without hydrolytic capacity

Because hypha-holding phagocytic cups structurally resemble tPCs, which display characteristics of both phagocytic cups and canonical phagosomes, we sought to determine the stage of phagocytic maturation reached by these compartments by following the recruitment of phagosomal maturation markers. As shown in **Figure 2 A**, hypha-holding tPCs formed in RAW cells presented with IgG-opsonized fixed hyphae were chiefly positive for the late phagosomal maturation markers Rab7 and Lamp-1 (**Figure 2 A ii)**, whereas early phagosomal markers were largely absent. Specifically, Rab5 was never detected at the phagocytic cup, and most cups were negative for PI(3)P (**Figure 2 A i**). Since hyphal engulfment is a continuous process, cup length can be used as a proxy for progression through engulfment and the associated phagosomal maturation timeline^21^. We therefore assessed phagosomal maturation kinetics by fitting logistic regression models that describe the probability of detecting each maturation marker as a function of cup length. Overall, the resulting fitted curves predict that as tPCs deepen to internalize longer segments of hyphae, they exchange PI(3)P for the late endo-lysosomal markers Rab7 and Lamp-1 (**Figure 2 B**). Furthermore, in accordance with another defining feature of phagosomal maturation, these tPCs fused with dextran loaded endo-lysosomes (**Figure 2 C i-ii**). The fitted logistic regression curve predicts that the frequency of endo-lysosomal fusion events increases as the cups deepen to engulf longer sections of the hyphal tips (**Figure 2 C iii**). However, unlike phagosomes, these cups remain open to the extracellular milieu. This became evident from their inability to retain fluid-phase cargo when endo-lysosomes were loaded with 10 kDa dextran (**Supplementary Figure 2 A**), in contrast to the retention observed with 70 kDa dextran (**Figure 2 C**). Importantly, hypha-holding tPCs formed during Dectin-1A mediated phagocytosis exhibited the same maturation and structural features of those formed through IgG-mediated phagocytosis (**Supplementary Figure 2 B-D**), therefore indicating that the tPCs characteristics described above are primarily dependant on the morphology of the target rather than the type of receptor involved in triggering phagocytosis.

**Figure 2.**
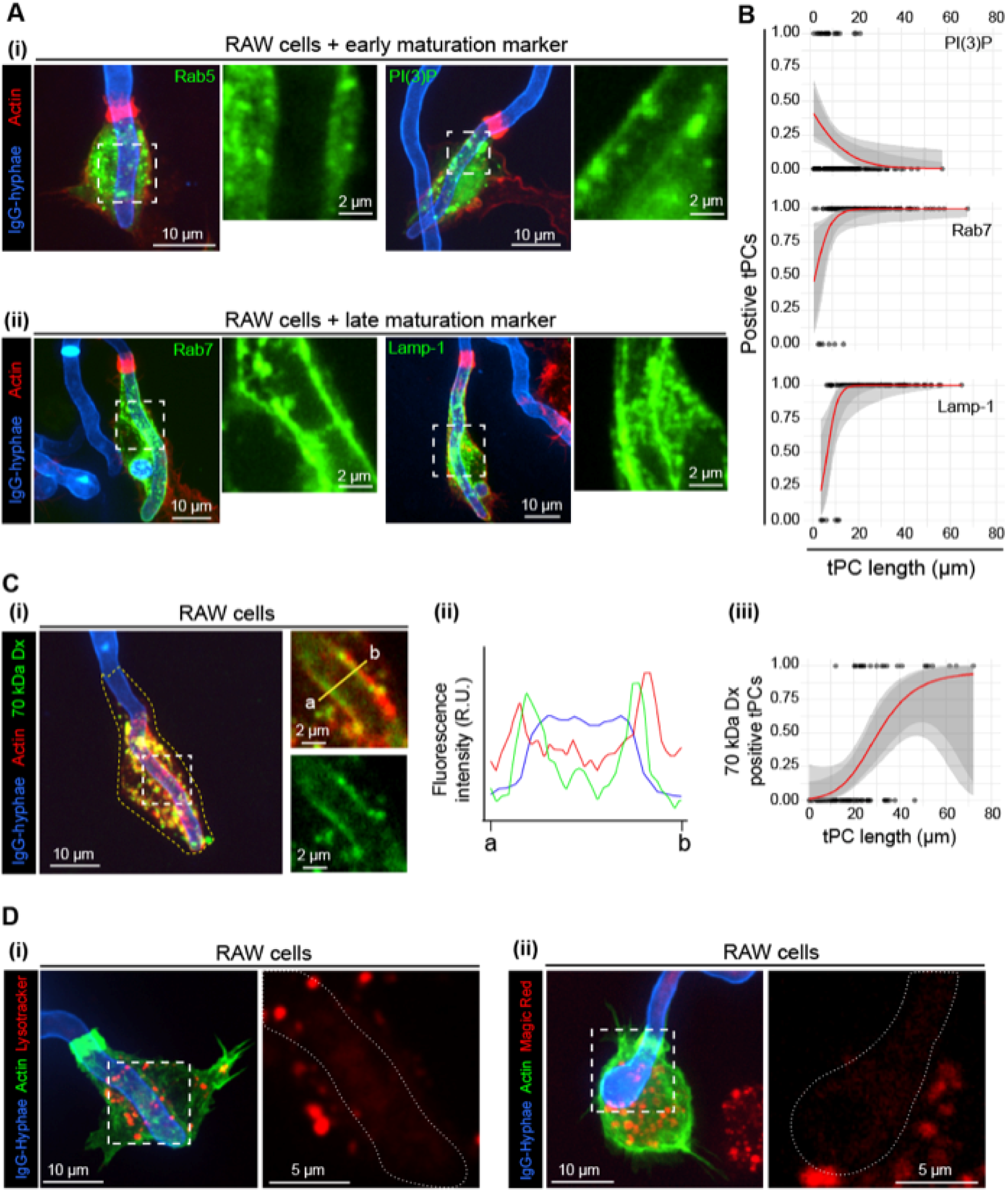
Grasps are tubular phagocytic cups that fuse with endosomal and lysosomal compartments but lack degradative capacity. (**A** and **B**) Acquisition of endosomal and lysosomal maturation markers by tubular phagocytic cups (tPCs). (**A**) RAW macrophages expressing EGFP-Rab5, GFP-2FYVE [PI(3)P biosensor], EGFP-Rab7, or LAMP-1-GFP (green) were incubated with IgG-hyphae (blue) for 1 hr, stained with phalloidin (red), and imaged by spinning-disk confocal microscopy (SDCM). Representative images and magnified views show the localization of the indicated maturation markers at hypha-holding tPCs. (**B**) Logistic regression models showing the probability of tPC acquisition of the indicated maturation markers [PI(3)P, LAMP-1, and Rab7] as a function of cup length. (**C**) Fluid-phase cargo accumulation within tPCs. RAW macrophages expressing LAMP-1-GFP (pseudocolored red) were pulse-labeled with tetramethylrhodamine (TMR)-conjugated 70-kDa dextran (70-kDa Dx, green) for 1 hr, chased for 2 hrs, incubated with IgG-hyphae as in (**A**), and imaged by SDCM. (**i**) Representative confocal images and magnified views show the distribution of LAMP-1 and 70-kDa Dx across a tPC cross-section. (**ii**) Representative fluorescence intensity profiles measured along the yellow line indicated in (**i**). (**iii**) Logistic regression model showing the probability of 70-kDa Dx accumulation within tPCs as a function of cup length. Dx = dextran. (**D**) Lysosomal acidity and proteolytic activity at tPCs. RAW macrophages expressing F-tractin-EGFP were incubated with (**i**) LysoTracker or (**ii**) Magic Red and subsequently challenged with IgG-hyphae as in (**A**). Representative SDCM images are shown.

Although hypha-holding tPCs acquired late phagosomal markers, their open nature prevented the development of key features of phagosome maturation. Accordingly, these compartments failed to accumulate the fluorescent probes LysoTracker™, which labels acidic compartments, and Magic Red™, which detects cathepsin activity, demonstrating an absence of luminal acidification and hydrolytic activity, respectively (**Figure 2 D**).

### Hypha-holding tPCs are sites of active ROS production

Since hydrolytic activity was not detected within the hypha-holding tPCs, we next examined whether these compartments produce ROS, which are a key antimicrobial mechanism deployed by phagocytes during phagocytosis ^32^. To assess whether ROS are locally produced within tPCs, we used the Nitroblue Tetrazolium (NBT) reduction assay that, in the presence of superoxide anion radicals (O_2_^·^) forms formazan, a purple precipitate that can be visualized by brightfield light microscopy ^33^. As shown in **Figure 3 A** and the corresponding quantitative data in **Figure 3 B**, tPCs formed by RAW cells in response to IgG-opsonized hyphae exhibited robust ROS accumulation that persist for at least 10 hrs. Consistent with a sustained ROS production, the cytosolic NOX2 subunit p67^phox^ was robustly recruited to both short and long hypha-holding tPCs (**Figure 3 C**). Importantly, similar results were observed in hypha-holding tPCs formed during Dectin-1A mediated phagocytosis (**Figure 3 D-F**), demonstrating that these open compartments sustain a localized and prolonged oxidative response, irrespective of the phagocytic receptor being engaged. This sustained ROS production contrasts sharply with the transient oxidative burst characteristic of canonical phagosomes, in which ROS generation typically persists for only ∼ 30 min ^34^.

**Figure 3.**
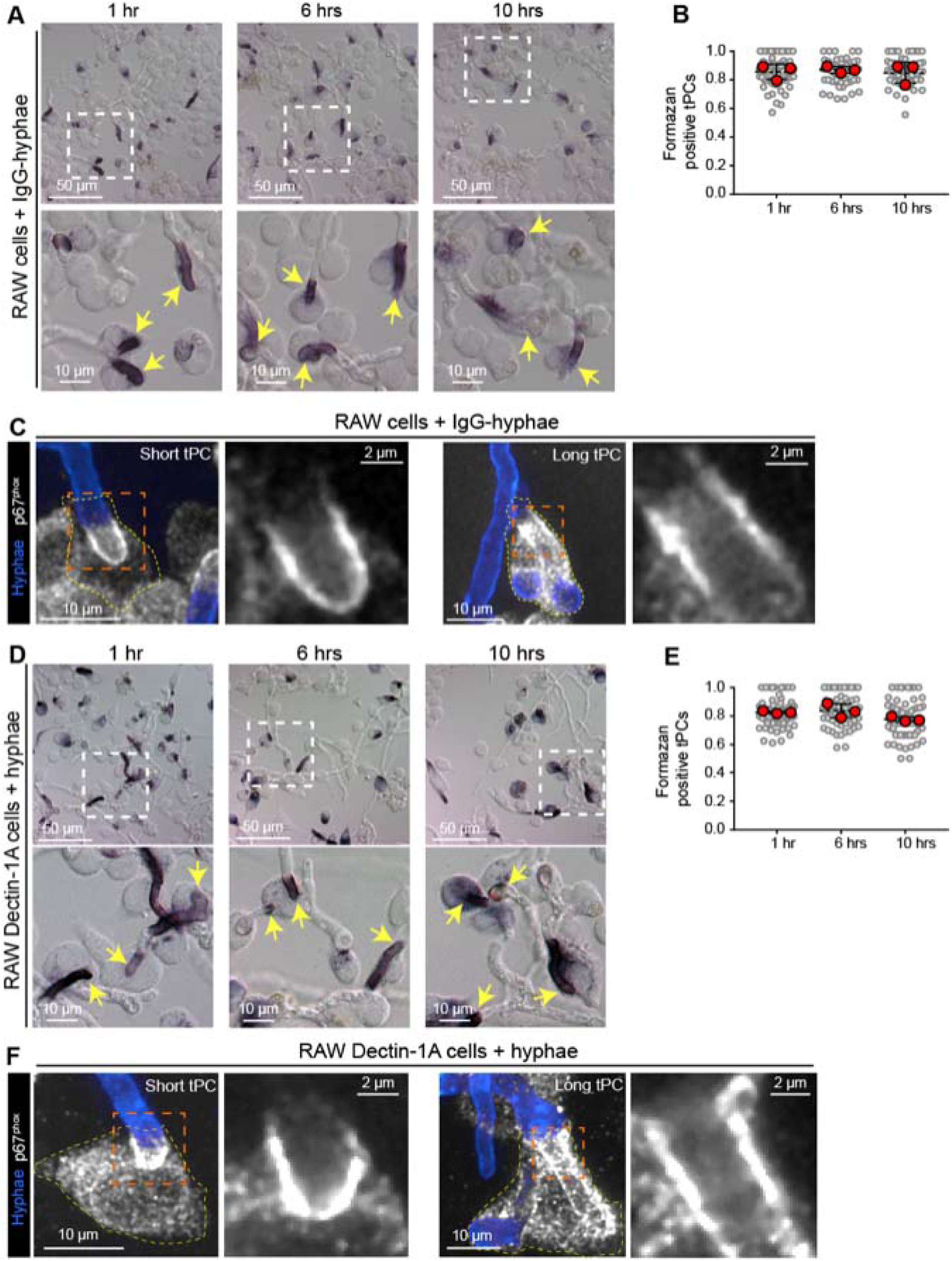
Prolonged ROS production and NOX2 machinery are detected at phagocytic cups in RAW macrophages. (**A-C**) Sustained ROS production and NOX2 recruitment at tPCs in RAW macrophages. (**A**) RAW macrophages were incubated with IgG-hyphae for the indicated times, followed by ROS detection by NBT assay and differential interference contrast (DIC) microscopy. Yellow arrows in the magnified views indicate formazan-positive tPCs. (**B**) Quantification of the proportion of formazan-positive tPCs. (**C**) RAW macrophages were incubated with IgG-hyphae (blue) for 1 hr, stained for p67^phox^ (gray), and imaged by spinning-disk confocal microscopy (SDCM). (**D-F**) Sustained ROS production and NOX2 recruitment at tPCs in RAW Dectin-1A macrophages. (**D**) RAW Dectin-1A macrophages were incubated with hyphae for the indicated times followed by ROS detection by NBT assay and DIC microscopy. (**E**) Quantification of the proportion of formazan-positive tPCs. (**F**) RAW Dectin-1A macrophages were incubated with hyphae labeled with calcofluor (blue) for 1 hr, stained against p67^phox^ (gray), and imaged by SDCM. (**B** and **E**) Gray points correspond to individual imaging fields and red points correspond to biological replicate means. Dashed horizontal lines indicate the overall mean of biological replicate means, and black error bars indicate SD. Statistical significance was determined by one-way ANOVA followed by Tukey’s multiple-comparisons test performed on biological replicate means (n = 3 independent experiments). ns (unlabeled), *P* > 0.05; \**P* < 0.05; \*\**P* < 0.01; \*\*\**P* < 0.001.

### Hyphal damage and growth control occurs in a ROS-dependent manner

Next, we sought to investigate whether the sustained ROS response observed within the hypha-holding tPCs exerts antifungal effects. Hyphal cell wall integrity is vital for fungal viability, as its disruption increases osmotic fragility and can ultimately result in cell lysis and death ^35^. Therefore, we assessed hyphal cell wall damage to determine whether sustained exposure to ROS within tPCs has an antifungal effect. To visualize the cell wall at sites of macrophage engagement, we used Calcofluor White (henceforth referred to as calcofluor), a fluorescent dye that binds chitin, a major component of the fungal cell wall. Calcofluor staining revealed a progressive loss of cell wall integrity in the tips of IgG-opsonized fixed hyphae entrapped within RAW tPCs. As shown in **Figure 4 A**, whereas 1 hr of macrophage-hyphal tip engagement resulted in no detectable effect to cell walls, hyphal tips entrapped for 6 hrs within tPCs exhibited extensive cell wall disruption. Consistent with this, time-lapse video microscopy monitoring live *A. fumigatus* hyphae revealed that capture of hyphal tips by macrophages within tPCs was associated with a pronounced reduction in apical growth, with elongation either slowing substantially or halting altogether (**Supplementary Video 3** and **Figure 4 B**, top panel). This growth-restrictive effect of tPCs was highly dependent on ROS, as macrophages lacking a functional NOX2 complex failed to effectively suppress hyphal elongation (**Supplementary Video 3, Figure 4 B**, lower panel, and **Figure 4 C**). Indeed, the capacity to halt hyphal apical growth was markedly impaired in RAW p22^phox^ KO cells, which are unable to assemble a functional NOX2 complex and therefore failed to generate ROS within hypha-holding tPCs (**Supplementary Figure 3 A**). Although RAW p22^phox^ KO cells retained the ability to capture hyphal tips, the entrapped apices frequently continued to elongate, ultimately piercing through the macrophage cells (**Supplementary Video 3** and **Figure 4 B**, lower panel; green trajectories). Importantly, similar results were obtained in RAW Dectin-1A cells phagocytosing non-opsonized hyphae (**Figure 4 D-F**, **Supplementary Video 4**, and **Supplementary Figure 3 B**), demonstrating that ROS-dependent hyphal damage and inhibition of apical growth are conserved features of tPCs that occur independently of the phagocytic receptor engaged.

**Figure 4.**
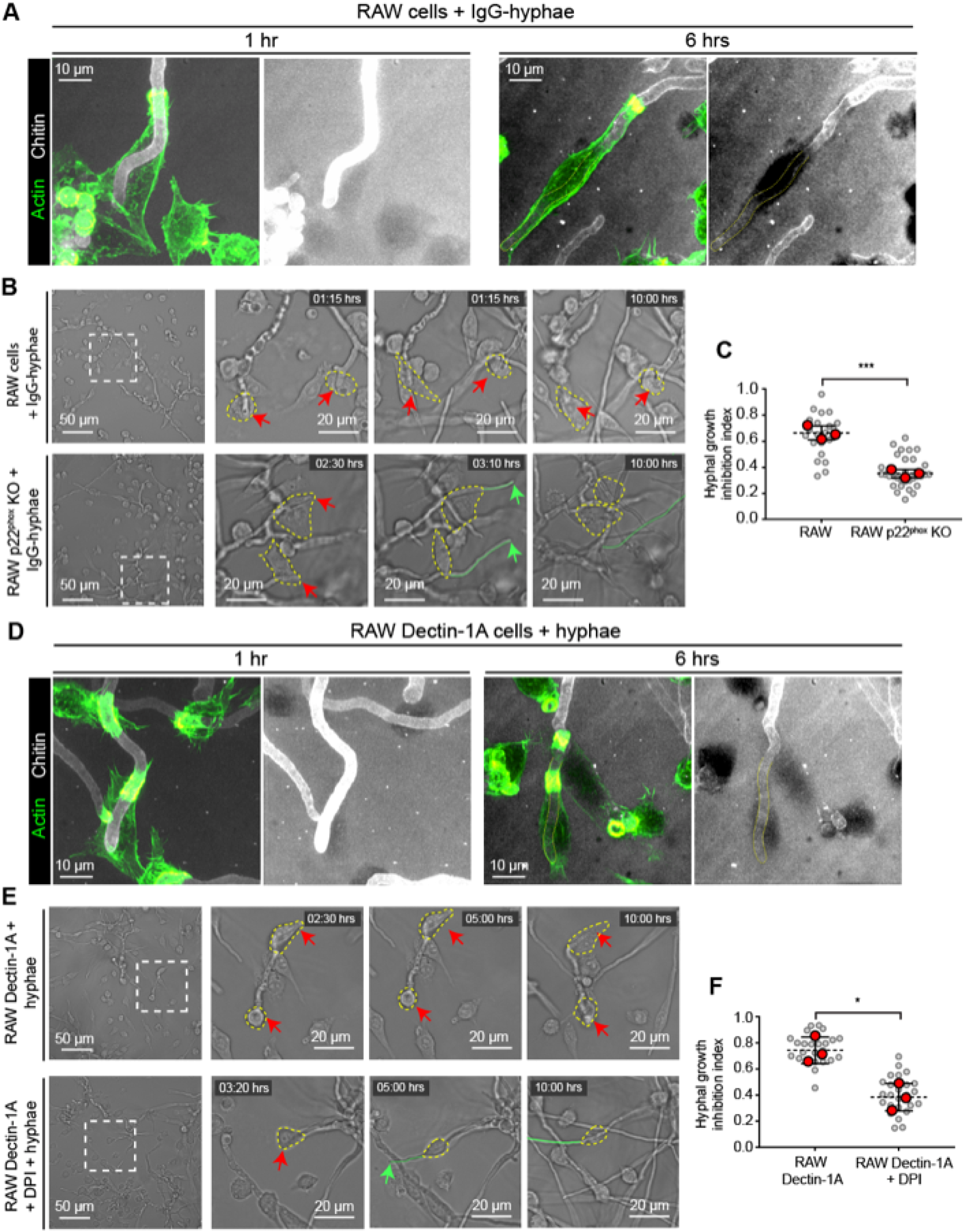
ROS production in phagocytic cups is associated with hyphal damage and apical growth control in RAW macrophages. (**A-C**) Hyphal damage and apical growth control by RAW macrophages. (**A**) RAW macrophages were incubated with IgG-hyphae (calcofluor stained following sample fixation, gray) for 1 or 6 hrs, stained with phalloidin (green), and imaged by SDCM. Representative images show merged projections (left) and the corresponding calcofluor channel alone (right). For the calcofluor-only images, signal intensity was linearly adjusted to enhance visualization of changes in hyphal tip cell wall staining. Yellow dotted line outlines region of hypha with depleted chitin staining. (**B**) RAW (top row) and RAW p22^phox^ KO (bottom row) macrophages were incubated with live IgG-hyphae and imaged by temperature-controlled brightfield time-lapse microscopy for 10 hrs. Images show representative stills from Videos S3. (**C**) Quantification of the hyphal apical growth inhibition index determined from the experiments described in (**B**). (**D-F**) Hyphal damage and apical growth control by RAW Dectin-1A macrophages. (**D**) RAW Dectin-1A macrophages were incubated with hyphae (calcofluor stained following sample fixation, gray) for 1 or 6 hrs, stained with phalloidin (green), and imaged by SDCM. Representative images show merged projections (left) and the corresponding calcofluor channel alone (right). For the calcofluor-only images, signal intensity was linearly adjusted to enhance visualization of changes in hyphal tip cell wall staining. Yellow dotted line outlines regions of hypha with depleted chitin staining. (**E**) RAW Dectin-1A (top row) and DPI treated Dectin-1A (bottom row) macrophages were incubated with live hyphae and imaged by temperature-controlled brightfield time-lapse microscopy for 10 hrs. Images show representative stills from Videos S4. (**F**) Quantification of the hyphal apical growth inhibition index determined from the experiments described in (**E**). (**B** and **E**) Macrophages engaging hyphal tips are outlined in yellow. Red arrows indicate hyphal tips retained within macrophage grasps, whereas green arrows indicate escaped hyphal tips. Green traces indicate subsequent hyphal growth. (**C** and **F**) Gray points correspond to individual imaging fields and red points correspond to biological replicate means. Dashed horizontal lines indicate the overall mean of biological replicate means, and black error bars indicate SD. Statistical significance was determined by an unpaired two-tailed t test performed on biological replicate means (n = 3 independent experiments). ns (unlabeled), *P* > 0.05; \**P* < 0.05; \*\**P* < 0.01; \*\*\**P* < 0.001.

### Primary human macrophages form tPCs that restrict hyphal growth

We next sought to validate our findings in primary human macrophages differentiated from peripheral blood monocytes (hMΦ). These cells express a broad repertoire of receptors for fungal recognition, including Dectin-1 ^36^. Accordingly, they did not require opsonization to engage and phagocytose live *A. fumigatus* hyphae within long-lasting tPCs (**Figure 5 A** and **Supplementary Video 5**).

**Figure 5.**
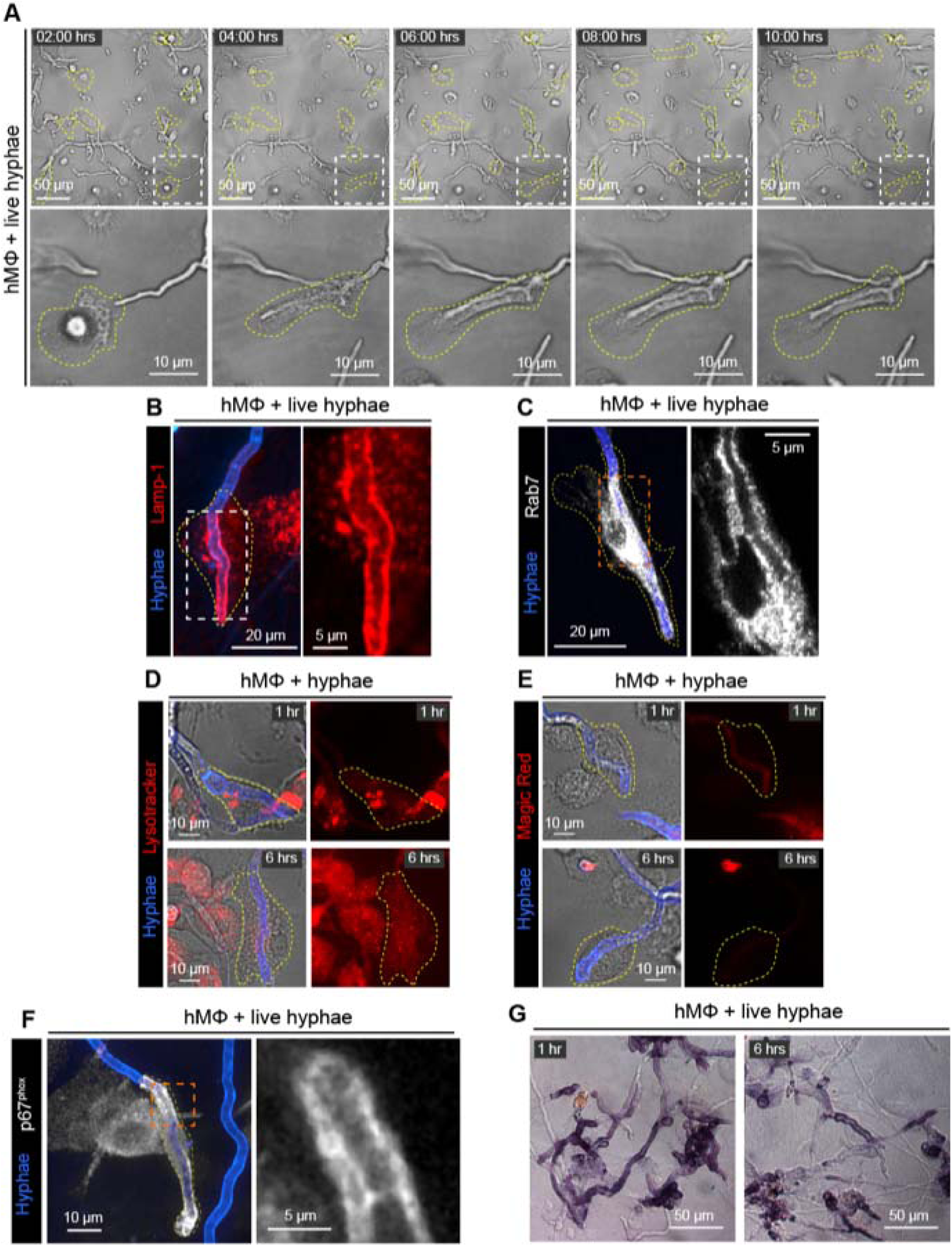
hMΦ form long-lasting phagocytic cups that acquire late phagosome maturation markers and sustain oxidative capacity but lack degradative capacity. (**A-C**) tPC formation and maturation in hMΦ. (**A**) hMΦ were incubated with live hyphae and imaged by temperature-controlled brightfield time-lapse microscopy for 10 hrs. Images show representative stills from Video S5. (**B-C**) Primary human macrophages were incubated with live hyphae for 1 hr, stained with calcofluor and against (**B**) LAMP-1 or (**C**) Rab7 following fixation, and imaged by SDCM. (**D** and **E**) Degradative capacity of hypha-holding tPCs in hMΦ. hMΦ were incubated with PFA-fixed hyphae for 1 or 6 hrs and loaded with (**D**) LysoTracker or (**E**) Magic Red. Representative SDCM images are shown. (**F** and **G**) Oxidative capacity of hypha-holding cups in hMΦ. (**F**) hMΦ were incubated with live hyphae (calcofluor-stained following fixation, blue) for 1 hr, stained against p67^phox^ (gray), and imaged by SDCM. Representative images and magnified views show p67^phox^ localization at phagocytic cups. (**G**) hMΦ were incubated with live hyphae for 1 or 6 hrs, followed by ROS detection by NBT assay and DIC microscopy.

As in the case of RAW and RAW Dectin-1A cells, tPCs formed in primary hMΦ acquired the late maturation markers Lamp-1 and Rab7 (**Figure 5 B, C**) but failed to acidify and develop hydrolytic activity (**Figure 5 D-E**). This contrasted with phagosomes containing *A. fumigatus* conidia and germlings or shorter hyphae which were both acidic and hydrolytic (**Supplementary Figure 3 C**). These hypha-holding tPCs were also positive for the NOX2 subunit p67^phox^ (**Figure 5 F**) and sustained ROS production for long periods (**Figure 5 G**). As shown for RAW and RAW Dectin-1A cells, the hyphae within these cups displayed extensive damage in their cell walls, as revealed by chitin staining (**Figure 6 A**). We then asked whether the cell wall damage was associated with osmolysis of hyphal apex cells. To this end, we challenged hMΦ with live *A. fumigatus* hyphae and employed the membrane impermeable vital dye Trypan Blue, which can permeate through the molecular sieves that gate tPCs ^19^. Following 6 hrs of engagement, Trypan Blue entered hyphal tips entrapped within tPCs, diffusing distally beyond the cup boundaries along the hyphal filament. This demonstrates the loss of plasma membrane integrity in cells at the hyphal apex (**Figure 6 B,** upper panel). As shown in the quantitative data of **Figure 6 C** (control plots), the proportion of hyphal damage increased drastically by 6 hours of engagement. Furthermore, consistent with what we showed for RAW and RAW Dectin-1A cells, the capacity of primary hMΦ to inflict hyphal damage was ROS-dependent, as treatment with the NOX2 inhibitor DPI prevented hyphal damage (**Figure 6 B**, **Figure 6 C** and **Supplementary Figure 3 D**), effectively precluding growth inhibition (**Figure 6 D-E** and **Supplementary Video 6**). Taken together, these findings demonstrate that hMΦ can capture hyphal tips within highly oxidative tPCs that damage the fungal cell wall, compromise apical cells integrity, and consequently halt hyphal growth.

**Figure 6.**
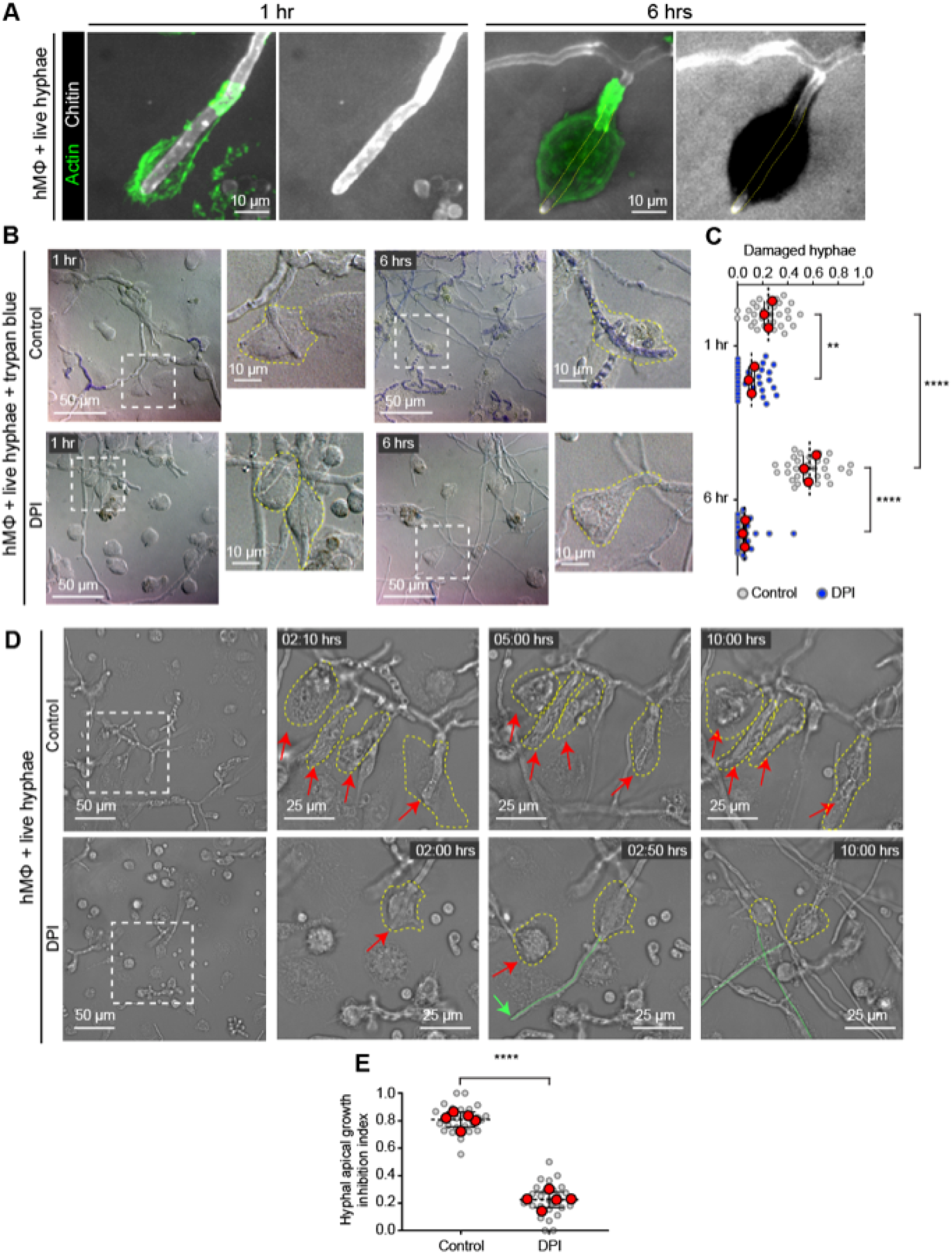
ROS accumulation in phagocytic cups is associated with hyphal damage and growth control in hMΦ. (**A-C**) Hyphal damage in hMΦ. (**A**) hMΦ were incubated with live hyphae (calcofluor stained following sample fixation, gray) for 1 or 6 hrs, stained with phalloidin (green), and imaged by SDCM. Representative images show merged projections (left) and the corresponding calcofluor channel alone (right). For the calcofluor-only images, signal intensity was linearly adjusted to enhance visualization of changes in hyphal tip cell wall staining. Yellow dotted lines outline regions of hyphae with depleted chitin staining. (**B**) hMΦ treated with DMSO (control; top row) or DPI (bottom row) were incubated with live hyphae for 1 or 6 hrs, followed by Trypan Blue staining to assess fungal plasma membrane damage. Representative images were acquired by DIC microscopy. (**C**) Quantification of the proportion of engaged hyphae exhibiting cytosolic Trypan Blue accumulation under conditions described in (**B**). (**D** and **E**) Hyphal growth control by hMΦ. (**D**) hMΦ (top row) and DPI-treated hMΦ (bottom row) were incubated with live hyphae and imaged by temperature-controlled brightfield time-lapse microscopy for 10 hrs. Images show representative stills from Video S6. Macrophages engaging hyphal tips are outlined in yellow. Red arrows indicate hyphal tips retained within macrophage grasps, whereas green arrows indicate escaped hyphal tips. Green traces indicate subsequent hyphal growth. (**E**) Quantification of the hyphal apical growth inhibition index determined from the experiments described in (**D**). (**C** and **E**) Gray or blue points correspond to individual imaging fields, and red points correspond to biological replicate means. Dashed horizontal lines indicate the overall mean of biological replicate means, and black error bars indicate SD. Statistical significance was determined using an unpaired two-tailed t test when comparing two means or one-way ANOVA followed by Tukey’s multiple-comparisons test when comparing > 2 means, performed on biological replicate means (n ≥ 3 independent experiments). ns (unlabeled), *P* > 0.05; \**P* < 0.05; \*\**P* < 0.01; \*\*\**P* < 0.001.

### PI(3,4)P_2_ production mediates ROS-dependent antifungal activity within tPCs

We next sought to determine the mechanisms underlying sustained ROS production at hypha-holding tPCs. Because PI(3)P is a key regulator of NOX2 activity and ROS production in canonical early phagosomes ^37^, and because we observed transient PI(3)P accumulation at the onset of hyphal phagocytosis (**Figure 2 A-B**), we investigated whether this phosphoinositide similarly contributes to the observed sustained ROS production within hypha-holding tPCs. To this end, we treated RAW cells with the class III PI3K inhibitor Vps34-IN prior to the onset of phagocytosis to block Vps34-dependent PI(3)P generation ^38^. Surprisingly, despite PI(3)P depletion on tPCs, robust ROS production persisted within these compartments (**Figure 7 A** and **B**). Although Vps34 inhibition produced a minor but statistically significant reduction in the proportion of formazan-enriched tPCs (**Figure 7 B ii**), the majority remained ROS positive, indicating that PI(3)P does not account for the bulk of ROS within these tPCs. Consistent with these findings, inhibition of the Vps34 had no detectable effect on ROS production in hypha-holding tPCs formed in RAW Dectin-1A cells (**Figure 7 C-D**).

**Figure 7.**
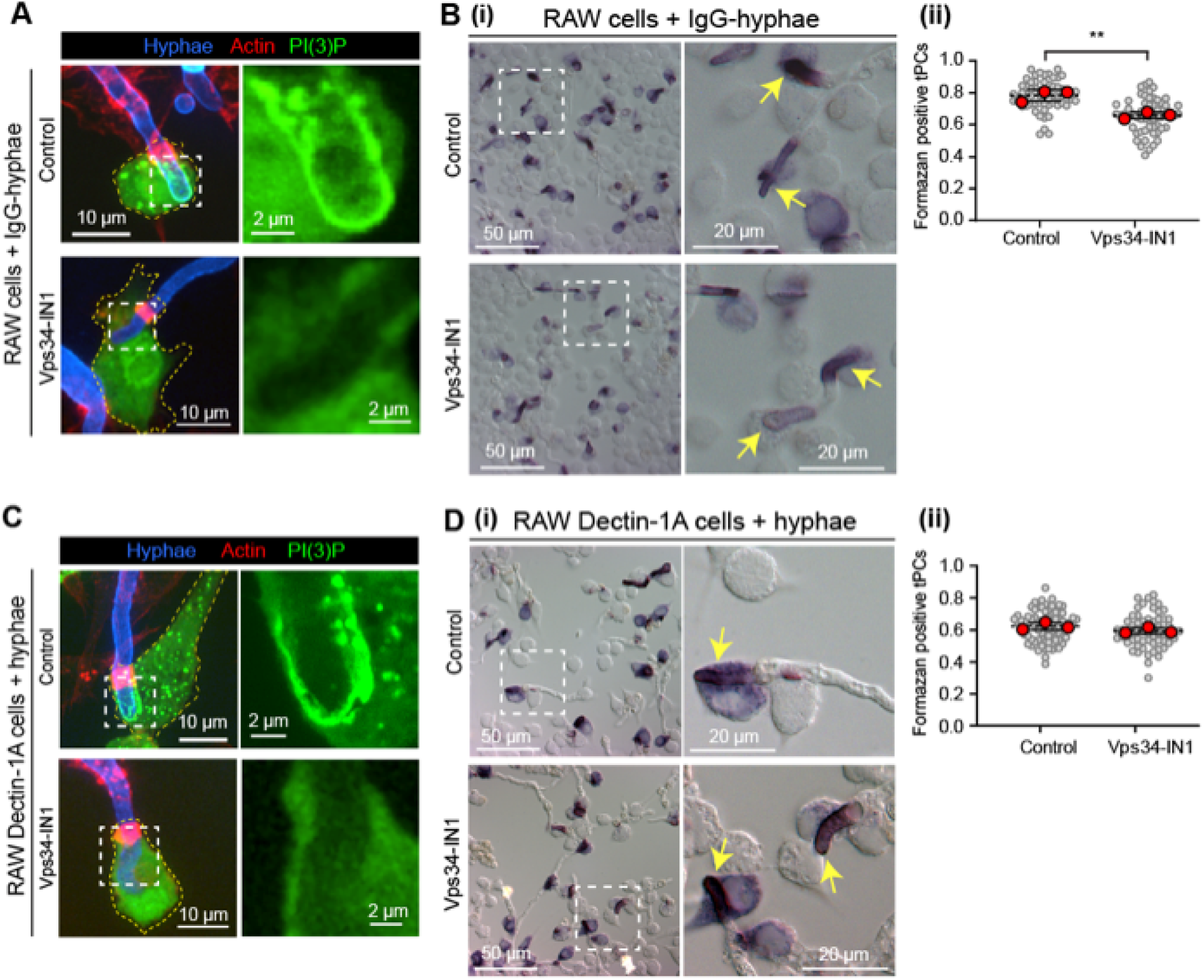
ROS production in phagocytic cups is not dependent on PI(3)P in RAW macrophages. (**A** and **B**) PI(3)P accumulation and ROS production at tPCs in RAW macrophages. (**A**) RAW macrophages expressing GFP-2FYVE [PI(3)P biosensor] treated with DMSO (top row) or Vps34-IN1 (bottom row) were incubated with IgG-hyphae (calcofluor-labeled, blue) for 1 hr, stained with phalloidin (red), and imaged by SDCM. (**B**) RAW macrophages treated with DMSO (top row) or Vps34-IN1 (bottom row) were incubated with IgG-hyphae followed by ROS detection by NBT assay. (**i**) Representative DIC micrographs. Yellow arrows in the magnified views indicate formazan-positive tPCs. (**ii**) Quantification of the proportion of formazan-positive tPCs. (**C** and **D**) PI(3)P accumulation and ROS production at tPCs in RAW Dectin-1A macrophages. (**C**) RAW Dectin-1A macrophages expressing GFP-2FYVE treated with DMSO (top row) or Vps34-IN1 (bottom row) were incubated with hyphae (calcofluor-labeled, blue) for 1 hr, stained with phalloidin (red), and imaged by SDCM. (D) RAW Dectin-1A macrophages treated with DMSO (top row) or Vps34-IN1 (bottom row) were incubated with hyphae followed by ROS detection by NBT assay. (**i**) Representative DIC micrographs. Yellow arrows in the magnified views indicate formazan-positive tPCs. (**ii**) Quantification of the proportion of formazan-positive tPCs. (**B ii** and **D ii**) Gray points correspond to individual imaging fields and red points correspond to biological replicate means. Dashed horizontal lines indicate the overall mean of biological replicate means, and black error bars indicate SD. Statistical significance was determined by an unpaired two-tailed t test performed on biological replicate means (n = 3 independent experiments). ns (unlabeled), *P* > 0.05; \**P* < 0.05; \*\**P* < 0.01; \*\*\**P* < 0.001.

The dissociation between ROS and PI(3)P occurrence at hypha-holding tPCs could also be inferred from the PI(3)P dynamics modelled in **Figure 2 B**, showing that while PI(3)P appears early during phagocytosis, it disappears as tPCs elongate. Yet, ROS remained persistently enriched across cups of all lengths (**Figure 3**). Together, these indicate that sustained ROS production and the resulting antifungal oxidative damage in hypha-holding tPCs are largely uncoupled from PI(3)P signaling.

We next examined PI(3,4)P_2_, another 3’-phosphoinositide generated during phagocytosis. PI(3,4)P_2_ is a relatively minor plasma membrane species that transiently accumulates at phagocytic cups during canonical phagocytosis and is typically lost following phagosome sealing ^39^. Moreover, PI(3,4)P_2_ has been associated with the regulation of NOX2 activity through structural and biochemical studies^40,41^. Thus, we assessed the occurrence of PI(3,4)P_2_ on hyphal-holding tPCs, by utilizing the recently developed and highly specific PI(3,4)P_2_ biosensor NES-EGFP-cPHx3 ^42^. As shown in **Figure 8 A** and **Supplementary Video 7**, PI(3,4)P_2_ was strongly enriched along the limiting membrane of hypha-holding tPCs, persisting for at least 6 hours.

**Figure 8.**
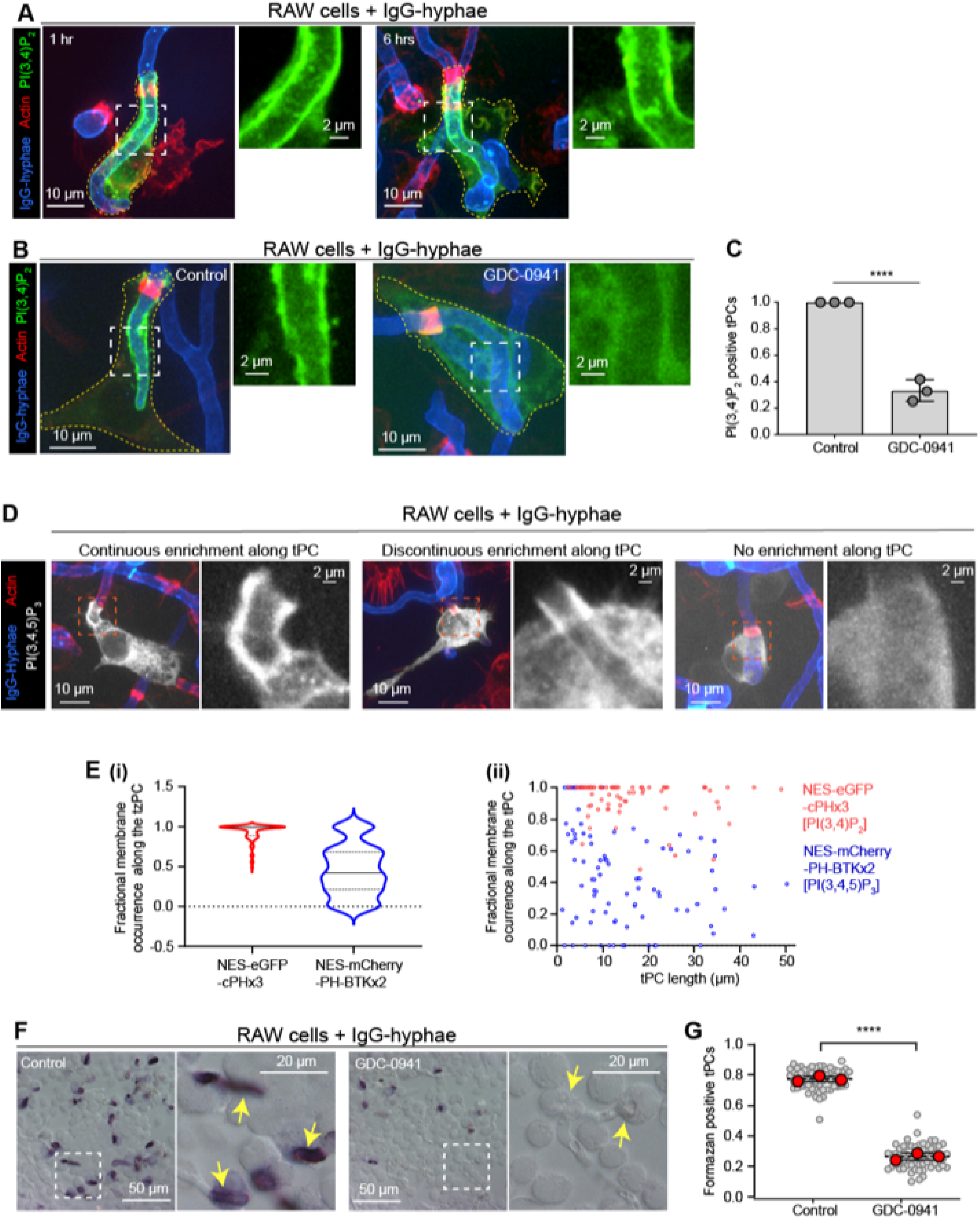
PI(3,4)P_2_ is enriched in phagocytic cups and explains ROS accumulation in RAW macrophages. (**A-C**) PI(3,4)P_2_ accumulation at tPCs in RAW macrophages. (**A**) RAW cells expressing NES-eGFP-cPHx3 [PI(3,4)P_2_ biosensor] were incubated with IgG-hyphae (calcofluor-labeled, blue) for 1 or 6 hrs, stained with phalloidin (red), and imaged by SDCM. Representative images are shown. (B) RAW cells expressing NES-eGFP-cPHx3 treated with DMSO (control) or GDC-0941 were incubated with IgG-hyphae (Calcofluor labelled, blue), stained and imaged as in (**A**). (**C**) Quantification of the proportion of PI(3,4)P_2_ -positive tPCs under the conditions described in (**B**). Gray points correspond to individual biological replicates. Black error bars indicate SD. (**D-E**) PI(3,4)P_2_ and PI(3,4,5)P_3_ enrichment at tPC membranes. (**D**) RAW macrophages expressing NES-mCherry-PH-BTKx2 [PI(3,4,5)P_3_ biosensor, pseudocoloured gray] were incubated with IgG-hyphae (calcofluor-labeled, blue) for 1 hr, stained with phalloidin (red), and imaged by SDCM. Representative images are shown. (**E**) Quantification of biosensor fractional occupancy at 1 hr tPCs. (**i**) Violin plots showing the distribution of fractional membrane positivity. (**ii**) Fractional membrane positivity of individual tPCs as a function of cup length. (**F-G**) Class I PI3K dependency of ROS production at tPCs. (**F**) RAW macrophages treated with DMSO (control) or GDC-0941 were incubated with IgG-hyphae followed by ROS detection by NBT assay. Representative DIC micrographs are shown. Yellow arrows in the magnified views indicate tPCs. (**G**) Quantification of the proportion of formazan-positive tPCs. Gray points correspond to individual imaging fields and red points correspond to biological replicate means. Dashed horizontal lines indicate the overall mean of biological replicate means, and black error bars indicate SD. (**C** and **G**) Statistical significance was determined by an unpaired two-tailed t test performed on biological replicate means (n = 3 independent experiments). ns (unlabeled), *P* > 0.05; \**P* < 0.05; \*\**P* < 0.01; \*\*\**P* < 0.001.

The dynamics of PI(3,4)P_2_ at the hypha-holding tPCs prompted us to investigate whether this phosphoinositide might be associated with the PI(3)P-independent ROS production. PI(3,4)P_2_ can be generated through different biosynthetic routes. Class II PI3Ks generate PI(3,4)P_2_ directly from the 3’ phosphorylation of PI(4)P ^43^. In addition, PI(3,4)P_2_ can be generated by a less characterized route involving the 4-phosphorylation of PI(3)P ^44^. Alternatively, the dephosphorylation of class I PI3K product, phosphatidylinositol 3,4,5-triphosphate [PI(3,4,5)P_3_] through 5’ phosphoinositide phosphatases has been shown to produce PI(3,4)P_2_ in the context of phagocytosis ^43^. Thus, we next sought to inhibit PI(3,4)P_2_ production by targeting these routes and assessing the resulting effects on ROS generation within hypha-holding tPCs.

Neither the inhibition of the class III PI3K with Vps34-IN1, nor the inhibition of class II PI3K with PITCOIN4 ^45^, interfered with the accumulation of PI(3,4)P_2_ or the production of ROS at hypha-holding tPCs (**Supplementary Figure 4 A-B** and **Figure 7 B**). In contrast, inhibition of class I PI3Ks with GDC-0941 ^46^ markedly depleted PI(3,4)P_2_ from hypha-holding tPCs (**Figure 8 B-C**). Together, these findings strongly suggest that the persistent pool of PI(3,4)P_2_ at hypha-holding tPCs is generated predominantly through the 5’ dephosphorylation of PI(3,4,5)P_3_, the product of class I PI3K activity. Several 5′ phosphatases are recruited during phagocytosis, with SHIP1 and 2 being logical candidates given their established role in generating PI(3,4)P_2_ from PI(3,4,5)P_3_^47,48^. However, the pharmacological inhibition of SHIP1 and 2 did not produce any detectable changes on PI(3,4)P_2_ dynamics at hypha-holding tPCs (data not shown). As previously reported for canonical phagocytosis ^39,49^, our observations point to functional redundancy among 5′ phosphatases and suggests that multiple enzymes contribute to sustaining PI(3,4)P_2_ production at hypha-holding tPCs.

Next, to assess the role of PI(3,4,5)P_3_ as a potential precursor of PI(3,4)P_2_, we monitored PI(3,4,5)P_3_ dynamics using the highly specific biosensor NES-mCherry-PH-BTKx2 ^50^. In contrast to PI(3,4)P_2_, which was continuously distributed along the limiting membrane of all hypha-holding tPCs (**Figure 8 A**), PI(3,4,5)P_3_ displayed marked heterogeneity in its localization. As shown in **Figure 8 D**, in elongated cups, PI(3,4,5)P_3_ distributed discontinuously and was preferentially enriched beneath the actin ring (middle panel), whereas continuous occurrence along the tPCs was only found in a subset of short cups (left panel). Moreover, a substantial fraction of hypha-holding tPCs lacked detectable PI(3,4,5)P_3_ signal altogether (right panel). This heterogeneity is captured in the quantitative analysis from **Figure 8 E i**, which measures the fractional membrane occupancy of each phosphoinositide biosensor along the limiting membrane of the cup and across tPC populations. Whereas PI(3,4)P_2_ exhibited a narrow distribution centered near complete membrane occupancy (fractional occupancy ≈ 1), PI(3,4,5)P_3_ displayed a much diffuse distribution, indicating considerable variability in its membrane occurrence. Stratification of fractional occupancy as a function of cup length further confirmed the persistent and widespread enrichment of PI(3,4)P_2_ along hypha-holding tPC membranes, whereas PI(3,4,5)P_3_ displayed substantial variability in membrane occupancy across these cups. Notably, PI(3,4,5)P_3_ was more consistently detected along the membrane of shorter tPCs (< 5 μm) but it became increasingly heterogeneous as cup length increased (**Figure 8 E ii**). These population-level measurements closely mirrored the localization patterns we observed by live-cell microscopy analysis (**Supplementary Videos 7** and **8**). Taken together, the class I PI3K-dependent PI(3,4)P_2_ enrichment and the contrasting spatiotemporal dynamics of PI(3,4)P_2_ and PI(3,4,5)P_3_ are consistent with a model in which transient PI(3,4,5)P_3_ production contributes to the generation of a more stable PI(3,4)P_2_ pool at hypha-holding tPCs.

Having established a class I PI3K-dependent pathway supporting persistent PI(3,4)P_2_ enrichment, we next asked whether disruption of this pathway affects sustained ROS production at hypha-holding tPCs. As shown in **Figure 8 F-G**, Inhibition of class I PI3K with GDC-0941 abolished ROS production in the majority of hypha-holding tPCs. This class I PI3K dependency was recapitulated in RAW Dectin-1A cells, where inhibition with GDC-0941 similarly depleted PI(3,4)P_2_ and substantially reduced ROS-positive tPCs (**Supplementary Figure 4 C-F**).

To determine whether this mechanism is conserved in hMΦ, we transfected cells with a synthetic mRNA encoding the PI(3,4)P_2_ biosensor NES-EGFP-cPHx3 and challenged them with live *A. fumigatus* hyphae. Consistent with our observations in RAW and RAW Dectin-1A cells, PI(3,4)P_2_ was robustly enriched along the limiting membrane of hMΦ hypha-holding tPCs, and this enrichment was abolished following treatment with the class I PI3K inhibitor GDC-0941 (**Figure 9 A**), indicating that PI(3,4)P_2_ synthesis depends on PI(3,4,5)P_3_ production. Notably, in these macrophages, tPCs generated substantially higher levels of ROS than RAW or RAW Dectin-1A cells, resulting in more intense formazan deposition, necessitating optimization of the NBT reduction assay for the proceeding quantitative analysis (**Figure 5 G**, see materials and methods for more details). Yet, our quantitative readout, based on the presence or absence of formazan, showed limited dynamic range. Thus, although GDC-0941 significantly reduced the proportion of formazan-positive cups, the effect size was modest using this metric **(Figure 9 B i, ii)**. We therefore quantified formazan pixel intensity within individual cups as a more sensitive measure of ROS production in hMΦ **(Figure 9 B iii**). This analysis revealed a markedly stronger reduction in cup-associated ROS following PI3K inhibition, further supporting a role for PI(3,4)P_2_ in sustaining NOX2 activity at hypha-holding tPCs. Therefore, we next asked whether the ROS-dependent fungicidal function of hypha-holding tPCs required PI(3,4)P_2_. To this end, we allowed hMΦ to form hyphal-holding tPCs and subsequently treated with them GDC-0941 for up to 10 hrs. As shown in **Figure 9 C** and **Supplementary Video 9**, this treatment caused a marked significant reduction of hyphal growth. Collectively, these findings confirm the link of class I PI3K activity with the persistent enrichment of PI(3,4)P_2_ and ROS production at hypha-holding tPCs with the ability of macrophages to contain *A. fumigatus* hyphal growth.

**Figure 9.**
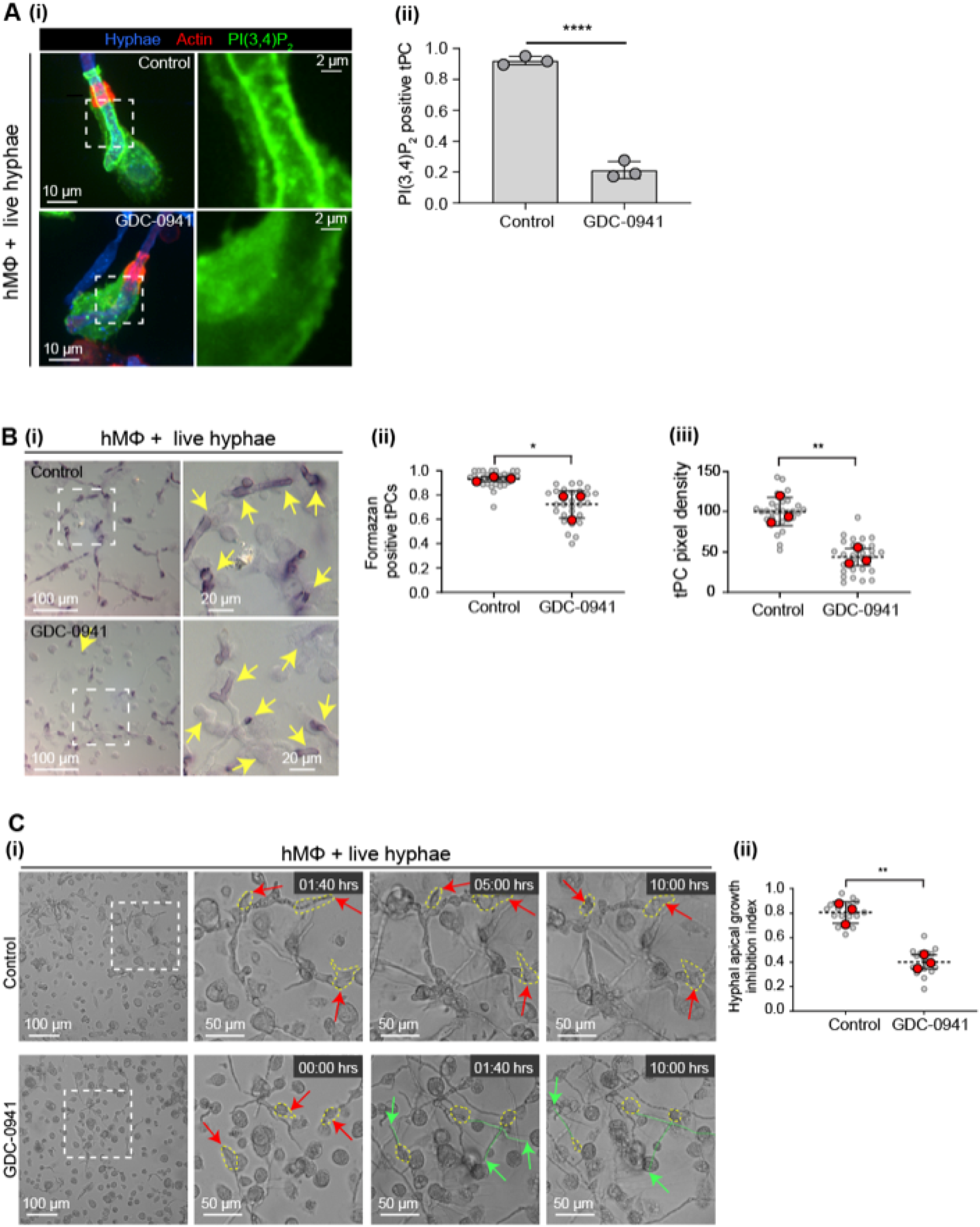
Class I PI3K activity is required for ROS accumulation in phagocytic cups and hyphal growth control in hMΦ. (A) Class I PI3K dependency of PI(3,4)P_2_ accumulation at tPCs in hMΦ. (**i**) hMΦ expressing mRNA encoding PI(3,4)P_2_ biosensor NES-eGFP-cPHx3 treated with DMSO (control) or GDC-0941 were incubated with live hyphae (calcofluor-labeled post-fixation, blue), stained with phalloidin (red), and imaged by SDCM. Representative images are shown. (**ii**) Quantification of the proportion of PI(3,4)P_2_ -positive tPCs under the conditions described in (**i**). Gray points correspond to individual biological replicates. Black error bars indicate SD. (B) Class I PI3K dependency of ROS production at tPCs in hMΦ. (i) hMΦ treated with DMSO (control) or GDC-0941 were incubated with live hyphae followed by ROS detection by NBT assay. Representative DIC micrographs are shown. Yellow arrows in the magnified views indicate tPCs. (**ii-iii**) Quantification of (**ii**) the proportion of formazan-positive tPCs and (**iii**) mean formazan pixel density within tPCs. (**B ii-iii** and **C ii**) Gray points correspond to individual imaging fields and red points correspond to biological replicate means. Dashed horizontal lines indicate the overall mean of biological replicate means, and black error bars indicate SD. (C) Class I PI3K dependency of hyphal growth control by hMΦ. (**i**) hMΦ treated with DMSO (control) or GDC-0941 were incubated with live hyphae and imaged by temperature-controlled brightfield time-lapse microscopy for 10 hrs. Images show representative stills from Videos S9. Macrophages engaging hyphal tips are outlined in yellow. Red arrows indicate hyphal tips retained within macrophage grasps, whereas green arrows indicate escaped hyphal tips. Green traces indicate subsequent hyphal growth. (**ii**) Quantification of the hyphal apical growth inhibition index determined from the experiments described in (**i**). (**A ii**, **B ii-iii** and **C ii**) Statistical significance was determined by an unpaired two-tailed t test performed on biological replicate means (n = 3 independent experiments). ns (unlabeled), *P* > 0.05; \**P* < 0.05; \*\**P* < 0.01; \*\*\**P* < 0.001.

## Discussion

It is now increasingly recognized that macrophages possess remarkable plasticity, enabling them to adapt their biology to the phagocytosis of targets whose dimensions challenge the conventional limits of engulfment ^12,15,19,22^. Such versatility may be critical in host defense against invasive filamentous fungi, such as *A. fumigatus*, where morphology is a determinant of pathogenicity ^2^. Yet, the prevailing paradigm assigns macrophages primarily to conidial clearance, whereas neutrophils are regarded as the dominant effector cells responsible for controlling hyphal growth ^1^. Here, we expand the current view of antifungal immunity by highlighting a previously underappreciated role for macrophages in the containment of *A. fumigatus* hyphae. In this cellular microbiology study, we show macrophages restrict *A. fumigatus* hyphal growth by forming persistent tPCs around hyphal tips, resulting in death of the key apical cells that control hyphal elongation. We proved that this antifungal activity depends on a sustained NOX2-derived ROS production, which requires persistent enrichment of PI(3,4)P_2_ at the cytosolic leaflet of the hypha-holding tPC membrane in both RAW cell lines and hMΦ.

Different from canonical phagocytic cups, hypha-holding tPCs acquire hallmarks of phagosomal maturation, marked by the sequential acquisition of PI(3)P, Rab7, and LAMP-1, and fuse with endo-lysosomal compartments despite remaining open compartments and maintaining a neutral luminal pH. This demonstrates that phagosomal maturation can proceed independently of phagosome sealing and acidification, both of which are generally considered *sine qua non* requirements for canonical phagosome maturation. Our findings are consistent with previous studies showing that elongated targets, including filamentous microorganisms and engineered rod-shaped particles, are internalized through long-lasting phagocytic cups that persist for extended periods before phagosome closure or, in some cases, remain incompletely sealed ^12,19,22,31^. Together, these observations suggest that persistent phagocytic cups represent a conserved adaptation of the phagocytic machinery for engaging targets whose geometry makes them difficult to engulf through canonical phagocytic mechanisms. In the case of fungal hyphae, including those of *A. fumigatus* or *C. albicans*, which can greatly exceed macrophage dimensions, this adaptation allows persistent interactions when complete target internalization is not possible ^12^. However, the antimicrobial properties of these structures appear to be highly context dependent. Whereas the tPCs described here act as antimicrobial compartments that capture hyphal tips and restrict *A. fumigatus* growth, prolonged cup persistence during the uptake of filamentous bacterial pathogens, including *L. pneumophila* and uropathogenic *Escherichia coli*, can delay phagosome completion and provide a window for phagocytic subversion and intracellular survival ^19,51^. Thus, persistent phagocytic cups may represent a broadly conserved response to elongated targets, but their biological outcome may be determined by the balance between host antimicrobial activity and pathogen adaptation.

Despite sharing several structural and molecular characteristics with the phagocytic cups formed around filamentous *L. pneumophila* ^19,31^, the tPCs that engage non-phagocytosable *A. fumigatus* hyphae display distinct maturation features. Specifically, we did not detect Rab5 recruitment to these compartments, despite the presence of Vps34-mediated PI(3)P production and Rab7 conversion, processes that are generally considered Rab5-dependent ^52^. These observations therefore raise the question of how maturation features are established in the apparent absence of detectable Rab5. One possible explanation is that Rab5 is recruited only transiently, with kinetics that fall below the temporal resolution of our imaging approach. This could be sufficient to support Vps34-mediated PI(3)P generation and Rab5-to-Rab7 conversion, thereby priming the phagocytic cup for fusion with endolysosomal compartments and accounting for the early acquisition of late-stage maturation markers. It also cannot be excluded that maturation of hypha-holding tPCs is mediated by LC3-associated phagocytosis (LAP), a form of non-canonical autophagy in which LC3 is conjugated to single-membrane phagosomes, thereby promoting lysosomal fusion and accelerating phagosome maturation ^53^. This possibility is particularly intriguing because LAP is triggered by ROS and has been implicated in the phagocytosis of *A. fumigatus* conidia ^54^. Notably, LAP has recently been shown to require the Rab5 isoform Rab5c ^55^, whereas conventional phagosome maturation primarily depends on Rab5a ^56^, the isoform examined in the present study. Thus, a LAP-dependent mechanism could explain the apparent lack of Rab5a despite the acquisition of phagosomal maturation features. Further studies will be required to determine whether LAP contributes to the maturation of hypha-holding tPCs.

The dynamics of the early phagosomal signaling lipid PI(3)P are markedly different in hypha-holding tPCs compared with phagocytic cups formed around filamentous bacteria. In the latter, PI(3)P is typically retained until near closure, or when the cup begins to acidify, triggering the dissociation of the Vps34 lipid kinase from the cup membrane ^31^. In contrast, hypha-holding tPCs rapidly lose PI(3)P, indicating distinct mechanisms regulating PI(3)P turnover at these structures. A plausible explanation is that hyphae and filamentous bacteria present markedly different physical and biochemical cues to phagocytes, including differences in diameter, rigidity, and surface molecular identity. These characteristics are likely to influence the signaling pathways engaged during engulfment thereby contributing to the distinct PI(3)P dynamics observed at hypha-holding tPCs.

Although PI(3)P is known to support ROS production by stabilizing the cytosolic NOX2 ternary complex (p40^phox^-p67^phox^-p47^phox^) in phagosomes^37^, the rapid loss of PI(3)P from hypha-holding tPCs, together with the observed insensitivity of ROS production to Vps34 inhibition, prompted us to conclude that sustained NOX2 activity in these structures is maintained through a PI(3)P-independent mechanism. Instead, we describe a sustained occurrence of PI(3,4)P_2_ at the cytosolic leaflet at the hypha-holding tPC membrane, which we found associated with continuous ROS production within the tPCs. The observed PI(3,4)P_2_ dynamics at tPCs is strikingly different from those reported for canonical phagocytosis, where this phosphoinositide occurs transiently at phagocytic cups to be rapidly depleted upon phagosome closure ^39^. These observations suggest that the open, hypha-holding tPC maintains a distinct phosphoinositide landscape that supports sustained NOX2 activity based on PI(3,4)P_2_. Although PI(3,4)P_2_ has been implicated in the recruitment, stabilization, and activation of NOX2 complex components at the plasma membrane, these biochemical and *in silico* studies did not address the role of PI(3,4)P_2_ within the cellular environment ^40,41^. Our findings expand this concept by identifying a role for PI(3,4)P_2_ in sustaining antimicrobial activity at hypha-holding tPCs through sustaining prolonged ROS production.

Beyond a role in sustaining ROS production, persistent PI(3,4)P_2_ accumulation at hypha-holding tPCs may also contribute to the maintenance of phagocytic cups through the recruitment of actin-regulatory effectors. Lamellipodin, a well-characterized PI(3,4)P_2_-binding protein, links phosphoinositide signaling to actin polymerization at sites of dynamic membrane remodeling and has been implicated in processes that support phagocytic cup formation ^39^. Similarly, TAPP1 has been linked to the regulation of actin cytoskeletal organization and membrane ruffling ^57^. Collectively, these observations suggest that sustained PI(3,4)P_2_ enrichment at tPCs serves, not only as a signaling platform associated with prolonged ROS production, but also as a membrane cue for the localized recruitment of actin-remodeling effectors. Such a mechanism could help preserve phagocytic cup architecture during the prolonged containment of fungal hyphae.

PI(3,4)P_2_ can arise through multiple biosynthetic routes ^43^. Although our data do not definitively identify its source at hypha-holding tPCs, several of our observations support that it is generated downstream of class I PI3K activity, most likely through the 5′-dephosphorylation of PI(3,4,5)P_3_. This interpretation is backed by the temporal dynamics of PI(3,4,5)P_3_ and PI(3,4)P_2_ at tPCs, as well as by the observed loss of PI(3,4)P_2_ following inhibition of class I PI3K. This, together with the persistent enrichment of PI(3,4)P_2_ at tPCs, suggest that continuous PI(3,4,5)P_3_ synthesis and turnover provide a sustained source of PI(3,4)P_2_ at the macrophage-hypha tPC interface.

Given that class I PI3Ks are activated downstream of multiple receptors implicated in fungal recognition, including phagocytic receptors, pattern-recognition receptors, and integrins, the sustained accumulation of PI(3,4)P_2_ likely reflects the continued integration of these signaling inputs during prolonged interactions of macrophages with *A. fumigatus* hyphae at the tPC interface. Persistent class I PI3K activity would be expected to continuously generate PI(3,4,5)P_3_, providing substrate for PI(3,4)P_2_ production. Because PI(3,4)P_2_ can be generated by multiple 5-phosphatases ^39,49^, pharmacological inhibition of SHIP1/2 did not prevent PI(3,4)P_2_ accumulation at hypha-holding tPCs. Together, these findings support a model in which sustained receptor-driven class I PI3K signaling maintains a local PI(3,4)P_2_-enriched membrane environment that promotes the assembly and/or stability of ROS-generating machinery, thereby enabling persistent ROS production at hypha-holding tPCs.

Our findings suggest that exposure of hyphal tips to this ROS response is the principal antifungal mechanism operating at tPCs. We propose that persistent oxidative stress at the hyphal apex disrupts Spitzenkörper function, thereby impairing the polarized trafficking of chitin synthases and other cell wall biosynthetic components required for apical growth ^30^. Consequently, localized cell wall assembly and remodeling may be compromised, resulting in reduced cell wall integrity and increased susceptibility of apical cells to osmotic lysis, ultimately impairing hyphal extension. However, damage to the apical chitin-rich cell wall was also observed in PFA-fixed hyphae, indicating that ROS can directly inflict oxidative damage to structural cell wall polysaccharides independently of fungal metabolic activities. Importantly, the latter is not mutually exclusive with ROS-mediated disruption of the apical Spitzenkörper-dependent growth machinery. Rather, both mechanisms may act in concert to compromise cell wall integrity and suppress hyphal extension at tPCs. Furthermore, we cannot exclude contributions from larger lysosomal hydrolases and antimicrobial protein assemblies that may be retained within the cup lumen, either because they exceed the size threshold for diffusion across the tPC interface, or through affinity-based interactions with the hyphal surface. Notably, chitotriosidase, a ∼ 50-kDa chitinase, remains active at near-neutral pH ^58^, and is therefore well suited to function within the non-acidified lumen of hypha-holding tPCs. Consequently, the local accumulation of lysosomal effectors within the tPC lumen may cooperate with ROS to compromise the fungal cell wall integrity and suppress hyphal growth.

Sustaining hypha-holding tPCs likely necessitates extensive focal exocytosis and mobilization of intracellular membrane reservoirs. Ongoing endolysosomal fusion may simultaneously support cup growth and replenish membrane-associated NOX2 components and lysosomal antimicrobial effectors. In this context, activation of lysosomal biogenesis programs would be expected to replenish the endolysosomal pools ^59^. Equally important, however, may be mechanisms that limit irreversible lysosome consumption by promoting membrane retrieval and recycling from the cup, thereby preserving cellular homeostasis and preventing exhaustion of the endolysosomal system ^60^. The prolonged generation of ROS within hyphal phagocytic cups presents an additional challenge, namely the potential for self-inflicted oxidative damage to host membranes through lipid peroxidation and membrane destabilization ^61^. Nevertheless, under our experimental conditions, we detected no evidence of cup membrane damage (data not shown), and the macrophage cytosol remained excluded from trypan blue following hyphal engagement, arguing against substantial compromise of tPC membrane integrity. These observations indicate that macrophages can effectively tolerate localized oxidative stress during prolonged interactions with fungal hyphae. Determining how these protective mechanisms cooperate to maintain the functionality of hypha-holding tPCs represents an important area for future investigation.

Together, our findings expand upon the current paradigm of antifungal immunity by demonstrating that macrophages engage directly with fungal hyphae and can exert antifungal activity through mechanisms distinct from canonical phagocytosis and intracellular killing. Thus, these results suggest that macrophages may serve as an important line of defense against hyphal growth, particularly in settings where neutrophil function is impaired or insufficient. Collectively, our data highlights a previously underappreciated role for macrophages and phagocytosis in host defense against *A. fumigatus* and more broadly, macrophage-mediated control of filamentous fungal pathogens.

## Materials and Methods

### Cell culture and reagents

RAW 264.7 murine macrophages (American Type Culture Collection; ATCC TIB-71™) were cultured in DMEM medium (Wisent Inc.) supplemented with 10% heat-inactivated fetal bovine serum (HI-FBS) (Gibco, Thermo Fisher Scientific). RAW cells stably expressing Dectin-1A, kindly provided by Dr. Nicolas Touret (University of Alberta, Edmonton Canada), were previously described (Lipinski et al., 2013) and maintained in the presence of 0.5 mg mL^−1^ geneticin (G418 sulfate; GIBCO, ThermoFisher Scientific). RAW H-2Kb p22^phox^ KO APOL7C::mCherry, herein referred to as RAW CRISPR-Cas9 p22^phox^ KO, is a CRISPR-Cas9-generated knockout clone lacking the p22^phox^ membrane-intrinsic subunit of NOX2 and contains a doxycyline-inducible APOL7C::mCherry construct. RAW CRISPR-Cas9 p22^phox^ KO cells were cultured in RPMI 1640 medium (Wisent Inc.) supplemented with 10% HI-FBS. All cell lines were maintained at 37°C in a humidified incubator with 5% CO2.

hMΦ were generated from peripheral blood monocytes isolated from healthy donors by density-gradient centrifugation using Lymphoprep^TM^ (STEMCELL Technologies). Monocytes were isolated by adherence and differentiated for 7-10 days in RPMI-1640 supplemented with 10% HI-FBS, 100 U mL^−1^ penicillin, 100 μg mL^−1^ streptomycin, 25 ng mL^−1^ <u>h</u>uman <u>M</u>acrophage <u>C</u>olony <u>S</u>timulating <u>F</u>actor (hM-CSF; Invitrogen, ThermoFisher Scientific). Cells were maintained at 37°C in a humidified incubator with 5% CO2. All procedures involving human samples were approved by the University of Toronto Health Sciences Research Ethics Board (Human Protocol #: 00043264).

### Fungal Strains and Culture Conditions

*A. fumigatus* strain UAMH 2978, a clinical isolate obtained from the UAMH Centre for Global Microfungal Biodiversity (Gage Research Institute, Canada), was grown on potato dextrose agar (PDA, BioShop Canada Inc.) at 28°C for 5-7 days. Conidia were harvested in potato dextrose broth (PDB, BioShop Canada Inc.) and 0.05% Tween 80 (PDB-T80) and filtered twice through three layers of sterile cheese cloth to remove conidiophores. Approximately 2 × 10^7^ conidia were added to 30 mL PDB-T80 and incubated at 37°C for 15 hrs to generate hyphae. Since conidial germination is asynchronous, hyphal samples include a wide range of target lengths (from 3 μm to more than 300 μm), and include conidia, swollen conidia and germlings that have not yet elongated to form hyphae.

For experiments using live hyphae, hyphae were washed three times with phosphate buffered saline (PBS) (1X) to remove residual Tween 80 and resuspended in the appropriate macrophage culture medium. For experiments using fixed hyphae, hyphae were fixed with 4% paraformaldehyde (PFA) for 1 hr at room temperature (RT), after which PFA was replaced with PBS and samples washed 3 times and stored at 4°C.

For opsonization with human IgG (human IgG, hIgG Sigma Aldrich), hyphae were pelleted by centrifugation at 10,000 g for 3 min at RT and washed three times with PBS (1X). Hyphae were then resuspended in 4 mg mL^−1^ h-IgG diluted in PBS (1X) and incubated on a rotating platform for 1 hr at RT or overnight at 4°C. Opsonized hyphae were subsequently pelleted as above and resuspended in the appropriate culture medium.

*Saccharomyces cerevisiae* INVSc1 was cultured according to the manufacturer’s instructions (Thermo Fisher Scientific). Overnight liquid cultures were harvested by centrifugation at 5000 g for 5 min, washed three times with PBS (1X), and fixed with 4% PFA for 30 min at RT. PFA was then replaced with PBS (1X), samples washed 3 times and stored at 4°C. h-IgG opsonization was performed as described above for *A. fumigatus*.

### Plasmids and DNA Transfection

The following plasmids were used in this work: EGFP-Rab5wt encoding wild type Rab5a isoform^62^; GFP-2FYVE^63^; EGFP-Rab7wt^62^; Lamp1-EGFP was purchased in Addgene (plasmid ID 16290); PM-GFP encoding the myristoylation/palmitoylation sequence of Lyn ^64^, F-tractin-eGFP (Addgene plasmid no. 58473); NES-eGFP-cPHx3 encoding the tandem carboxy-terminal PH (cPH) domains of TAPP1^42^ were purchased in Addgene (plasmid ID 116855), NES-GFP-PH-BTKx2 and NES-mCherry-PH-BTKx2 encoding the two tandem Bruton’s Tyrosine Kinase (BTK) PH domains^50^ were purchased in Addgene plasmid ID 183654 and 183654).

For transfections, RAW cells were seeded into plastic multi-well plates to 60–70% confluency and transfected using Lipofectamine™ LTX reagent with PLUS™ reagent (Thermo Fisher Scientific) according to the manufacturer’s instructions. At 14 hrs post-transfection, RAW cells were washed once with PBS (1X), detached by scrapping, and seeded onto glass coverslips to the desired confluency (40-60%). Phagocytosis assays (described below) were performed 6-12 hrs after reseeding.

### Customized mRNA Synthesis and Transfection of hMΦ

The mRNA encoding the PI(3,4)P2 biosensor NES-eGFP-cPHx3 was developed and synthesized via Northern RNA (Canada). Briefly, the plasmid codifying for the NES-eGFP-cPHx3 was used as a template for *in vitro* transcription of mRNA. The mRNA was co-transcriptionally capped using Co Cap A, and all uridine bases were substituted with N1-methylpseudouridine to improve mRNA stability, reduce cellular response to synthetic mRNA and enhance the expression of the encoded protein. Purified mRNA was resuspended in water (pH 7) at a concentration of 0.98 mg mL^−1^.

For mRNA transfections, peripheral blood monocytes isolated from healthy human donors were seeded onto 18 mm glass coverslips at a density of 1 × 10^6^ cells per coverslip and differentiated into macrophages as described above. Subsequently, macrophages were transfected with the NES-eGFP-cPHx3 mRNA using Lipofectamine^TM^ MessengerMax^TM^ (Thermo Fisher Scientific) according to the manufacturer’s instructions. Cells were incubated for 24 hrs following transfection before phagocytosis assays (described below) were performed.

### Phagocytosis Assays

*A. fumigatus* hyphae were either stained with Calcofluor White (calcofluor, Sigma Aldrich) before macrophage exposure, stained post-fixation, or left unstained, as indicated. For pre-staining, hyphae were pelleted at 10,000 g for 3 min at RT, resuspended in calcofluor diluted 1:100 in PBS (1X), and incubated for 5 min at RT on a rotating platform to achieve even staining of the hyphal cell wall. Hyphae were then washed three times with PBS, resuspended in the appropriate macrophage culture medium, and added to macrophages at a ratio of two fungal particles per macrophage. Hyphae were allowed to settle onto macrophages for 5 min at 37°C, followed by incubation at 37°C for the indicated times. Samples were subsequently fixed with 4% PFA for 15 min at RT.

For post-fixation calcofluor staining, hyphae were supplied to macrophages unstained and then samples were permeabilized with 0.1% Triton X-100 (BioShop Canada Inc.) for 15 min and stained with calcofluor (1:100 in PBS, 1X) for 5 min at RT on a Belly Dancer orbital shaker. Samples were then washed five times with PBS (1X) to remove residual calcofluor.

For phagocytosis of *S. cerevisiae* yeast, macrophages were equilibrated at 15°C for 10 min and subsequently challenged with yeast at a ratio of 20:1 yeast per macrophage. Samples were then centrifuged at 300 g for 5 min at 15°C, after which the medium was replaced with fresh culture medium, and cells were incubated at 37°C for the indicated times. Samples were then processed for ROS detection using the NBT assay and fixed with 4% PFA.

Where indicated, tPCs were visualized by staining actin with phalloidin-Alexa Fluor 555 (1:400 in PBS, 1X) (Invitrogen, Thermo Fisher Scientific) as per the manufacturer’s instructions. Following any additional staining procedures, coverslips were mounted using fluorescence mounting medium (Aligent Dako, Aligent Technologies) for microscopy analysis.

### Immunofluorescence

The following primary antibodies were used: mouse anti-human Lamp-1 monoclonal antibody (clone H4A3; Developmental Studies Hybridoma Bank), anti-NCF2 (p67^phox^) rabbit polyclonal antibody (cat. # PA5-37323; Invitrogen, Thermo Fisher Scientific), anti-Rab7 D95F2 (Cell Signaling, 9367S) Alexa Fluor 555 donkey anti-rabbit IgG (cat. # A31572; Life Technologies, Thermo Fisher Scientific) and Alexa Fluor 488 goat anti-mouse IgG (cat. # A31556; Invitrogen, Thermo Fisher Scientific).

For immunolabelling of endogenous Lamp-1, PFA-fixed samples were washed three times with PBS (1X) and permeabilized with methanol at −20°C for 10 min. Samples were blocked with 2.5% bovine serum albumin (BSA) diluted in PBS (1X) for 30 min at RT and incubated with anti-Lamp 1 antibodies (1:100) for 1 hour at RT. Samples were then washed three times with PBS (1X) and incubated with the corresponding secondary antibody (1:1500) for 1.5 hours at RT, followed by three PBS (1X) washes and mounted as described in previous sections. For immunolabelling of endogenous p67^phox^, PFA-fixed samples were washed three times with PBS (1X), permeabilized with 0.1% Triton X-100 for 15 min at RT, and blocked with 3% BSA in PBS (1X) for 30 min at RT. Samples were incubated with anti-p67^phox^ antibodies (1:100) overnight at 4 °C, washed three times with PBS (1X), and incubated with the corresponding secondary antibody (1:1000) for 2 hours at RT. Samples were then washed three times with PBS (1X) and mounted as described above. For immunolabelling of endogenous Rab7, PFA-fixed samples were washed three times with PBS (1X), permeabilized with 20µM digitonin () for 10 min at RT, and blocked with 10% skim milk in PBS (1X) for 1 hr at RT. Samples were incubated with anti-Rab7 antibodies (1:50) overnight at 4 °C, washed three times with PBS (1X), and incubated with the corresponding secondary antibody (1:100) for 1.5 hours at RT. Samples were then washed three times with PBS (1X) and mounted as described above.

### Fluorescence-based Labeling of Endolysosomal Compartments

For preloading endolysosomal compartments, a pulse/chase experiment was performed. Briefly, RAW cells expressing Lamp1-EGFP or F-tractin-EGFP were incubated with 0.1 mg mL^−1^ Alexa Fluor^TM^ 647-conjugated 10 kDa Dextran (Invitrogen, Thermo Fisher Scientific) or tetramethylrhoadamine (TMR)-conjugated 70 kDa Dextran (Invitrogen, Thermo Fisher Scientific) in culture medium for 1 hr at 37°C. Cells were subsequently washed three times with PBS (1X) and chased for 2 hrs at 37°C in culture medium before phagocytosis assays were performed. For labeling of acidic compartments, macrophages were incubated with 1 μM LysoTracker™ Deep Red (Invitrogen, Thermo Fisher Scientific) during the final hour of exposure to *A. fumigatus*.

To assess degradative capacity, macrophages were incubated with Magic Red Cathepsin L Assay Kit (Immunochemistry Technologies LLC) during the final 15 min of *A. fumigatus* exposure, according to the manufacturer’s instructions. Following labelling, cells were washed once with PBS (1X) and imaged live in complete culture medium lacking phenol red.

### Detection of ROS with Nitroblue Tetrazolium Assay

Following exposure to hyphae for the indicated times, RAW cells were incubated with 0.5 mg mL^−1^ Nitroblue Tetrazolium (NBT; BioShop Canada Inc.) for 30 min to detect ROS production. Cells were subsequently washed three times with PBS (1X), fixed with 4% PFA for 15 min, washed an additional three times with PBS (1X), and mounted as described above.

To adjust for higher levels of ROS production in hMΦ, NBT assays were performed by incubating cells with 0.5 mg mL^−1^ NBT for 10 min to avoid saturation of formazan accumulation within phagocytic compartments. For experiments involving pharmacological inhibitors, the respective treatments were maintained throughout NBT incubation period. Cells were analyzed using Differential Interference contrast (DIC) imaging as described below.

### Assessment of Hyphal Damage by Trypan Blue Staining

To assess hyphal cell wall damage, RAW or hMΦ were incubated with *A. fumigatus* hyphae for the indicated times. Samples were fixed with 4% PFA, permeabilized with 0.1% Triton X-100 for 15 min andwashed three times with PBS (1X). Hyphae were then stained with alcofluor (1:1000 in PBS, 1X) to reveal differences in chitin staining along the hyphal filament. Triton X-100 permeabilization was performed to ensure uniform access of calcofluor to the hyphal cell wall. F-actin was stained using phalloidin-Alexa Fluor 555 and mounted as described above.

To assess hyphal plasma membrane damage, hMΦ exposed to live hyphae were stained with the membrane-impermeable dye Trypan Blue (Invitrogen, Thermo Fisher Scientific). Trypan Blue accumulation within the hyphal cytosol was used as a proxy for compromised fungal plasma membrane integrity, as the dye can traverse the fungal cell wall but enters the cytosol when plasma membrane integrity is lost^65^. Briefly, samples were washed twice with PBS (1X) and incubated for 5 min at RT with 0.2% Trypan Blue diluted 1:1 in culture medium. Samples were immediately fixed with 4% PFA, gently washed once with PBS (1X), and mounted as described above.

### Treatment with pharmacological inhibitors

For inhibition of NADPH oxidase activity, RAW and hMΦ were pretreated with 10 mM diphenylene iodonium (DPI; Sigma-Aldrich) or DMSO as vehicle control (BioShop Canada Inc.) for 30 min prior to the commencement of phagocytosis assays, and treatments maintained throughout the duration of the experiments.

For inhibition of Vps34, RAW cells were pretreated with 1 mM Vps34-IN1 (Selleck Chemical) or DMSO for 1 hr, followed by phagocytosis in the continued presence of the respective treatments. Treatment after the onset of phagocytosis was to avoid interfering with the development of the tPCs.

For inhibition of Class I or Class II PI3K, RAW or RAW Dectin-1A cells were incubated with hyphae for 1 hr, followed by treatment with 0.5mM GDC-09450 (EMD Millipore) for Class I PI3K inhibition or 50 mM of PITCOIN4 for Class II PI3K inhibition, or DMSO for an additional 2 hrs. Treatments were maintained during subsequent NBT assays or analysis of phosphoinositide biosensor distribution.

To functionally validate PITCOIN4 activity, we assessed inhibition of clathrin-mediated endocytosis by the internalization of transferrin^66^. Briefly, RAW cells were incubated with 10 mM Pitstop2 (Abcam), a known inhibitor of clathrin-mediated endocytosis, 50 mM of PITCOIN4 or DMSO for 40 min, followed by incubation at 4°C for 30 min and then 5 min at 37°C in the presence of 25 mg/mL transferrin conjugated to Alexa Fluor 546 (Thermo Fisher Scientific). This was followed by two washes with PBS (1X) and two washes with 1.5 M acetic acid, followed by fixation as described above.

For hMΦ, a shorter treatment protocol with GDC-0941 was used to minimize differences in tPCs length between conditions. hMΦ were incubated with hyphae for 20 min, followed by treatment with 5 mM GDC-0941 or DMSO for 10 min prior to NBT assays or analysis of phosphoinositide biosensor distribution. For brightfield time-lapse imaging, hMΦ were incubated with live hyphae for 20 min followed by treatment with 5 mM GDC-0941 or DMSO, which was maintained throughout the remaining imaging period.

### Scanning Electron Microscopy

Following phagocytosis assays, RAW cells were fixed with 2% glutaraldehyde in 0.1 M sodium cacodylate buffer (pH 7.4) for 2 hrs at RT. Samples were washed three times in 0.1 M sodium cacodylate buffer for 5 min each and post-fixed with 1% OsO_4_ for 1 hr at RT. Samples were then washed three times with 0.1 M sodium cacodylate buffer, incubated with 1% tannic acid for 30 min at RT, and treated with 1% OsO_4_ for an additional 30 min. Samples were washed three times with double-distilled water for 10 min each, Subsequently, samples were sequentially dehydrated by incubation with increasing concentrations of ethanol (30%, 50%, 70%, 95%, 100% v/v) and the specimens were transferred to a critical point dryer (Leica Critical Point Dryer EM CPD300). Dry samples were mounted onto aluminum SEM stubs and transferred to a Osmium plasma coater (Filgen OPC60A / OPC-EVO). Surface imaging was performed using a Hitachi S-530 scanning electron microscope (Hitachi S530). Images were acquired and processed using Quartz PCI software (Quartz Imaging Corporation).

### Confocal, brightfield and DIC imaging and processing

Spinning disk confocal microscopy (SDCM) was performed using a Quorum Technologies WaveFX spinning disk confocal microscope equipped with a Yokogawa CSU-X1 spinning disk unit, Spectral Laser Merge Module (406, 491, 561, and 643 nm), and Hamamatsu EMCCD and Orca R2 cameras. Images were acquired using 40× 1.3 NA or 63× 1.4 NA oil-immersion objectives, with acquisition controlled by MetaMorph software (Molecular Devices).

For live-cell imaging, cells onto glass coverslips were enclosed in a Leiden chamber and immersed in live cell imaging solution (Molecular Probes™, Thermo Fisher Scientific) supplemented with 10% FBS and mounted in a microscope-mounted environmental chamber pre-warmed at 37°C. Where indicated, confocal imaging was performed using a Leica Stellaris 5 laser scanning confocal microscope (LSCM) equipped with a 406-nm violet laser, tunable white-light laser (485–685 nm), HyD detectors, and 40× 1.30 NA or 63× 1.4 NA oil-immersion objectives. Image acquisition was controlled using Leica Application Suite X (LAS X) software. DIC imaging for visualization of NBT-derived formazan and Trypan Blue staining was performed using a Zeiss Axioplan 2 (Zeiss) upright microscope equipped with an AxioCam HRc colour camera. Image acquisition was controlled using AxioVision software.

Long-term brightfield time-lapse imaging was performed using an Etaluma LS720 live-cell microscope (Etaluma) housed within a cell incubator maintained at 37 °C and 5% CO2. Cells were maintained in the appropriate culture medium, lacking phenol red for the duration of imaging. Time-lapse sequences were acquired using LumaView software.

For quantitative analyses of tPCs, structures were imaged as encountered within the field of view and were not selected based on cup morphology, length, marker recruitment, or other phenotypic features. Image processing and quantitative analyses were performed using Fiji^67^ or Volocity software (Quorum Technologies Inc.). Where applicable, image adjustments were performed uniformly and without altering the quantitative relationship between image elements. Figures and time-lapse sequences were prepared using Adobe Illustrator and Adobe Photoshop (Adobe).

### Quantification of formazan accumulation via image analysis

Formazan accumulation within tPCs was quantified by mean gray value using Fiji. Images were assessed for pixel saturation and overexposure using the HiLo lookup table and histogram tools in Fiji prior to quantitative analysis. Briefly, individual tPCs regions were manually outlined, and the mean gray value within each region was measured. Background mean gray values were calculated separately for each image and subtracted from the corresponding measurements on tPCs. Because increased formazan deposition results in lower gray values in transmitted-light images, background-corrected values were multiplied by −1 for graphical representation when comparing formazan accumulation across macrophage–hypha incubation times. For experiments comparing pharmacological treatments with vehicle controls, background-corrected values were normalized to the corresponding control condition.

### Logistic Regression Analysis

To model the probability of maturation marker acquisition as a function of tPCs length, logistic regression analysis was performed. Briefly, individual tPCs were classified as marker positive (1) or marker-negative (0), with tPC length used as the continuous predictor variable. To account for potential non-linear relationships between tPC length and marker acquisition, polynomial logistic regression models of increasing degree were evaluated, and the optimal polynomial degree was selected based on the Akaike Information Criterion (AIC). The resulting best-fitting model was used to estimate the probability of marker acquisition as a function of tPC length. Models were generated from pooled data from three independent experiments with at least 15 tPCs analyzed per biological replicate. Regression curves are shown in red, with dark and light grey regions representing the 95% and 99% confidence intervals, respectively.

### Quantification of fractional membrane occurrence at tPCs

Confocal image stacks were merged in Volocity, and the membrane of individual tPCs was manually traced. The total length of the cup membrane and the length of the membrane occurrence for the indicated markers were measured. Fractional membrane occurrence was calculated as the length of marker-positive cup membrane divided by the total traced cup membrane length. A value of 0 represents no detectable marker recruitment to the cup membrane, whereas a value of 1 represents marker positivity along the entire cup membrane.

### Statistical Analysis

Data is presented as mean ± standard deviation of at least three independent experiments, unless otherwise indicated. Statistical analysis was carried out using the Prism 9.0.2 software (GraphPad, La Jollam). Data was assumed to be normally distributed. Comparisons between two conditions were performed using an unpaired two-tailed Student’s *t* test. Comparisons among multiple conditions were performed using ordinary one-way ANOVA followed by Tukey’s multiple comparisons test t, as indicated. Statistical analyses were performed on biological replicate means. *P <* 0.05 was considered statistically significant. Statistical tests are specified in the corresponding figure legends.

## Author Contributions

Serene Moussaoui (SM) and Mauricio Roberto Terebiznik (MRT) are responsible for the overall design of this study. SM and Maria Cecilia Gimenez (MCG) designed and carried out experiments, data analysis, and interpretation. Amanda Rebouças Paixão (ARP), Melie Boulianne (MB), Amir Kanbar (AK), and Abdu Okhunjonov (AO) carried out experiments and data analysis. Federico Martinez (FM) and Micaela Cabrini (MC) performed data analysis. Carlos Alberto Merino (CAM) developed the R script and implemented the logistic regression analyses. Durga Acharya (DA) carried out the sample preparation for SEM analysis. SM and MRT wrote the manuscript. MRT and MCG reviewed and edited the manuscript. All authors gave approval to the final version of the manuscript.

## Acknowledgements

We thank Dr. Emily Rosowski (Clemson University, USA) for insightful discussions about the project. We thank Durga Acharya and Bruno Chue from the Centre for Microscopy and Bioanalysis (CMB) at the University of Toronto Scarborough for their invaluable support and advice with electron and fluorescence microscopy. We thank Gabriel Fabiano and Aliza Khaitin for technical support and troubleshooting during the experimental setup. We thank Dr. Jonathan Canton (University of Calgary, Edmonton Canada) for kindly sharing the g RAW CRISPR-Cas9 p22^phox^ KO, Dr. Sergey Plotnikov (University of Toronto) for gifting the F-tractin-eGFP constructs, and Dr. Marc Nazaré and Volker Haucke (Leibniz-Forschungsinstitut für Molekulare Pharmakologie, Berlin, Germany) for kindly providing the PITCOIN4.

This work was supported by the Natural Sciences and Engineering Research Council (NSERC) discovery grants RGPIN-2024-05569 to MRT. APR was a recipient of the UTSC postdoctoral fellowship. SM is a recipient of an NSERC PhD Student Research Award. AK was a recipient of an NSERC Undergraduate Student Research Award (USRA). AO was a recipient of the University of Toronto Excellence Awards (UTEA). MC was a recipient of a Mitacs Globalink Research Award (GRA). The CMB facility is funded by a Canada Foundation for Innovation (CFI) grant (#493864).

## Declaration of generative AI and AI-assisted technologies in the manuscript preparation process

During the preparation of this work, the author(s) used Microsoft 365 Copilot to refine scientific writing and improve the clarity of presentation. The author(s) reviewed and edited the output as needed and take full responsibility for the content of the published article.

## Declaration of interests

The authors declare no competing interests.

## Notes

### Competing Interest Statement

The authors have declared no competing interest.

