## Supplementary Figures and Captions for "3′ Plasma Membrane Phosphoinositides Sustain ROS Production at Persistent Non-canonical Phagocytic Cups that Promote Macrophage Control of *Aspergillus fumigatus* Hyphae"

**Supplementary Figures and Legends**

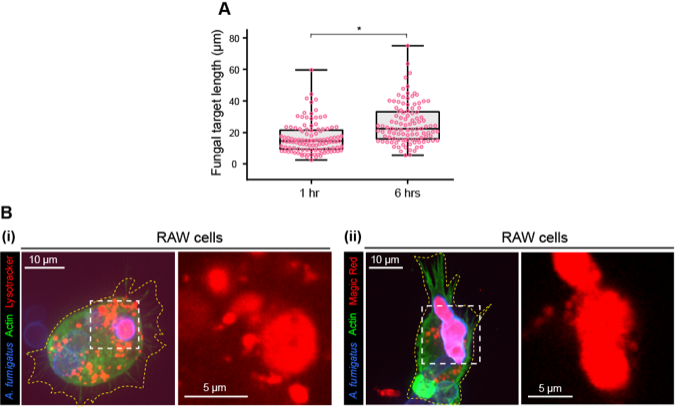

**Supplementary Figure 1.** **Characterization of *A. fumigatus* containing phagosomes in RAW macrophages**

(**A**) Length distribution of sealed phagosomes. RAW macrophages were incubated with IgG-hyphae (calcofluor-labeled, blue) for 1 or 6 hrs. Box plots show the distribution of phagosome lengths. Individual phagosomes were pooled from three independent experiments. Statistical significance was determined by an unpaired two-tailed t test performed on biological replicate means (n = 3 independent experiments). ns (unlabeled), *P* > 0.05; **P* < 0.05; ***P* < 0.01; ****P* < 0.001.

(**B**) Acidity and proteolytic activity of sealed phagosomes in RAW macrophages. RAW macrophages were incubated with IgG-hyphae (calcofluor-labeled, blue) for 1 hr and loaded with (**i**) LysoTracker or (**ii**) Magic Red. Representative SDCM images are shown.

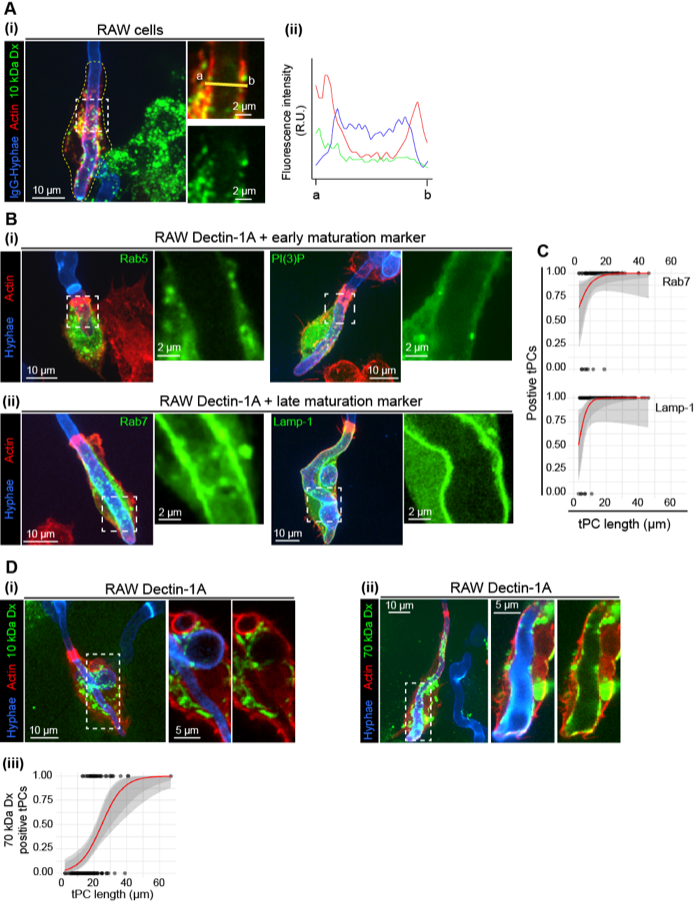

**Supplementary Figure 2.** **Validation of tPC characteristics in RAW and RAW Dectin-1A cells**

(**A**) Low molecular-weight fluid-phase cargo within RAW macrophages that form tPCs. RAW macrophages expressing LAMP-1-GFP (pseudocolored red) were pulse-labeled with Alexa Fluor 647-conjugated 10-kDa dextran, chased for 2 hrs, incubated with IgG-hyphae (calcofluor-labeled, blue) for 1 hr, and imaged by SDCM. (**i**) Representative confocal images and magnified views show the distribution of LAMP-1 and dextran across a tPC cross-section. (**ii**) Representative fluorescence intensity profiles measured along the yellow line indicated in (**i**). Dx = dextran.

(**B** and **C**) Acquisition of endosomal and lysosomal maturation markers by tPCs in RAW Dectin-1A macrophages. (**B**) RAW Dectin-1A macrophages expressing EGFP-Rab5, GFP-2FYVE [PI(3)P biosensor], EGFP-Rab7, or LAMP-1-GFP were incubated with hyphae (calcofluor-labeled, blue) for 1 hr, stained with phalloidin (red), and imaged by SDCM. Representative images and magnified views show the localization of the indicated maturation markers at hypha-holding tPCs. (**C**) Logistic regression models showing the probability of tPC acquisition of the indicated maturation markers [LAMP-1 and Rab7] as a function of cup length.

(**D**) Fluid-phase cargo accumulation within tPCs in RAW Dectin-1A macrophages. RAW Dectin-1A macrophages expressing F-tractin-eGFP were pulse-labeled with (**i**) Alexa Fluor 647-conjugated 10-kDa dextran or (**ii**) TMR-conjugated 70-kDa dextran, chased for 2 hrs, and challenged with hyphae (calcofluor-labeled, blue) for 1 h, and imaged by SDCM. Representative SDCM images are shown. (**iii**) Logistic regression model showing the probability of 70-kDa Dx accumulation within tPCs as a function of cup length.

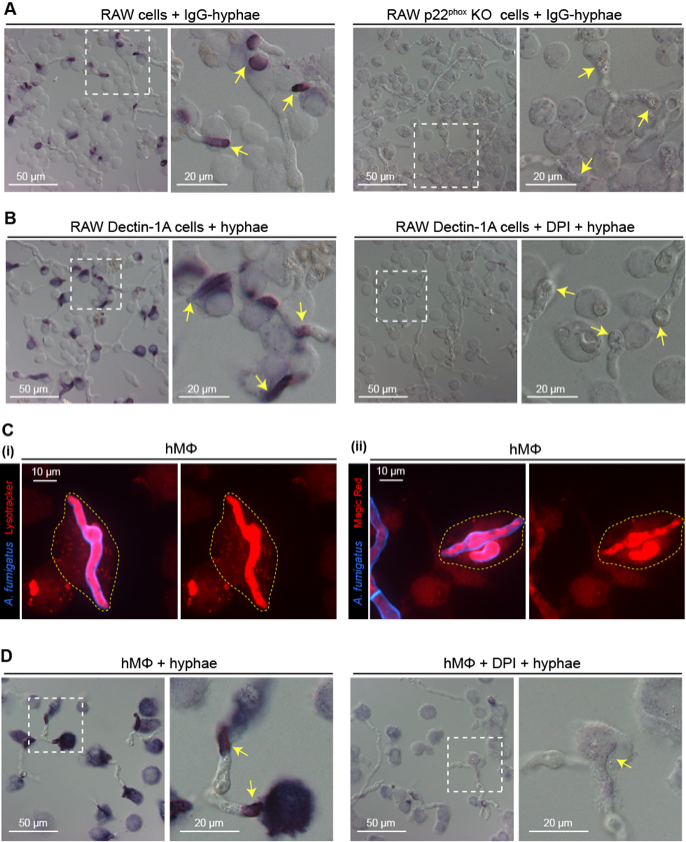

**Supplementary Figure 3.** **Validation of genetic and pharmacological inhibition of ROS production at phagocytic cups**

(**A-B, D**) ROS production at phagocytic cups following genetic or pharmacological inhibition of NOX2 activity. (**A**) RAW and RAW p22^phox^ KO macrophages were incubated with IgG-hyphae for 1 hr, followed by ROS detection by NBT assay. (**B**) RAW Dectin-1A macrophages and DPI-treated RAW Dectin-1A macrophages were incubated with hyphae for 1 hr, followed by ROS detection by NBT assay. (**D**) hMΦ and DPI-treated hMΦ were incubated with hyphae for 1 hr, followed by ROS detection by NBT assay. (**A-B, D**) Representative DIC micrographs are shown. Yellow arrows in the magnified views indicate phagocytic cups.

(**C**) Acidity and proteolytic activity of sealed phagosomes in hMΦ. hMΦ were incubated with fixed hyphae for 1 hr and loaded with (**i**) LysoTracker or (**ii**) Magic Red. Representative SDCM images are shown.

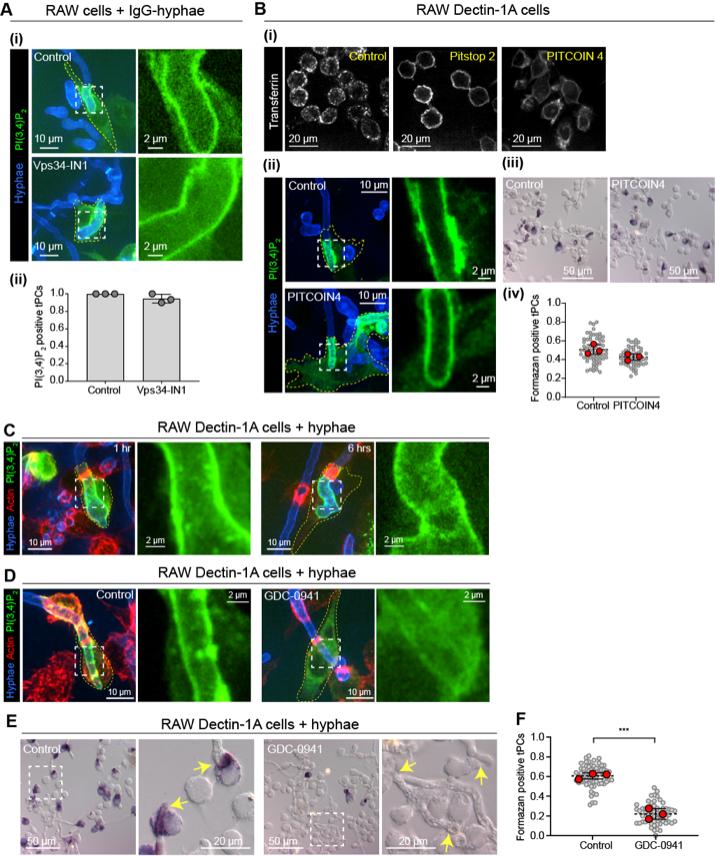

**Supplementary Figure 4.** **PI(3,4)P_2_ accumulation and ROS production at tPCs require Class I PI3K activity in RAW Dectin-1A cells**

(**A**) Class III PI3K dependency of PI(3,4)P_2_ accumulation at tPCs in RAW macrophages. (**i**) RAW macrophages expressing NES-eGFP-cPHx3 [PI(3,4)P_2_ biosensor] treated wiwth DMSO (control) or Vps34-IN1 were incubated with IgG-hyphae (calcofluor-labeled, blue) for 1 hr and imaged by SDCM. (**ii**) Quantification of the proportion of PI(3,4)P_2_ -positive tPCs in RAW macrophages treated with DMSO (control) or Vps34-IN1. Gray points correspond to individual biological replicates. Black error bars indicate SD.

(**B**) Class II PI3K dependency of PI(3,4)P_2_ accumulation and ROS production in RAW Dectin-1A macrophages. (**i**) RAW Dectin-1A macrophages were incubated with Alexa Fluor 546-conjugated transferrin in the presence of DMSO (control), Pitstop 2 or PITCOIN4 and imaged by SDCM. (**ii**) RAW Dectin-1A macrophages expressing NES-eGFP-cPHx3 [PI(3,4)P_2_ biosensor] were incubated with hyphae (calcofluor-labeled, blue) for 1 hr following treatment with DMSO (control) or PITCOIN4 and imaged by SDCM. (**iii**) RAW Dectin-1A macrophages treated with DMSO (control) or PITCOIN4 were incubated with hyphae followed by ROS detection by NBT assay. (**iv**) Quantification of the proportion of formazan-positive tPCs.

(**C-F**) Class I PI3K dependency of PI(3,4)P_2_ accumulation and ROS production at tPCs in RAW Dectin-1A macrophages. (**C**) RAW Dectin-1A macrophages expressing NES-eGFP-cPHx3 were incubated with hyphae (calcofluor-labeled, blue) for 1 or 6 hrs, stained with phalloidin (red), and imaged by SDCM. Representative images are shown. (**D**) RAW Dectin-1A macrophages expressing NES-eGFP-cPHx3 treated with DMSO (control) or GDC-0941 were incubated with hyphae (calcofluor-labeled, blue) and imaged by SDCM. Representative images are shown. (**E**) RAW Dectin-1A macrophages treated with DMSO (control) or GDC-0941 were incubated with hyphae followed by ROS detection by NBT assay. Representative DIC micrographs are shown. (**F**). Quantification of the proportion of formazan-positive tPCs within tPCs for conditions described in (**E**).

(**B iv** and **F**) Gray points correspond to individual imaging fields and red points correspond to biological replicate means. Dashed horizontal lines indicate the overall mean of biological replicate means, and black error bars indicate SD.

(**A ii**, **B iv** and **F**) Statistical significance was determined by an unpaired two-tailed t test performed on biological replicate means (n = 3 independent experiments). ns (unlabeled), *P* > 0.05; **P* < 0.05; ***P* < 0.01; ****P* < 0.001.

**Supplementary Video Captions**

**Supplementary Video 1.** Representative brightfield time-lapse microscopy of RAW macrophages engaging IgG-opsonized hyphae.

**Supplementary Video 2.** Representative brightfield time-lapse microscopy of RAW Dectin-1A macrophages engaging hyphae.

**Supplementary Video 3.** Representative brightfield time-lapse microscopy comparing hyphal growth control by RAW and RAW p22^phox^ KO macrophages.

**Supplementary Video 4.** Representative brightfield time-lapse microscopy comparing hyphal growth control by DMSO control and DPI-treated RAW Dectin-1A macrophages.

**Supplementary Video 5.** Representative brightfield time-lapse microscopy showing tPC formation and sustenance in hMΦ.

**Supplementary Video 6.** Representative brightfield time-lapse microscopy comparing hyphal growth control by DMSO control and DPI-treated hMΦ.

**Supplementary Video 7.** Representative SDCM time-lapse imaging of PI(3,4)P_2_ accumulation at tPCs in RAW macrophages expressing NES-eGFP-cPHx3.

**Supplementary Video 8.** Representative SDCM time-lapse imaging of PI(3,4,5)P_3_ accumulation at tPCs in RAW macrophages expressing NES-mCherry-PH-BTKx2.

**Supplementary Video 9.** Representative brightfield time-lapse microscopy comparing hyphal growth control by DMSO control and GDC-0941-treated hMΦ.
